# Chemogenetic excitation and play promote functional recovery in juvenile spinal cord injury

**DOI:** 10.64898/2026.09.24.754164

**Authors:** Rupert D. Smit, Thomas J. Campion, Ajay Pal, Joshua D. Ross, Jacquelynn Rajavong, Huiling Ke, Gavin T. Koma, Ismail Safak, Zenub M. Abouzid, Jose Paz Amaya, Michel Lemay, Andrew J. Spence, George M. Smith

**Affiliations:** Department of Neural Sciences, Center for Neural Development and Repair, Lewis Katz School of Medicine at Temple University, Philadelphia, 19140, USA; Department of Bioengineering, Temple University, College of Engineering, Philadelphia, 19140, USA; Appraiseye Inc., Babraham Research Campus, CB22 3FH, Cambridge, UK

**Keywords:** spinal cord injury, paediatrics, chemogenetics, corticospinal tract, neuroplasticity, rehabilitation

## Abstract

Functional recovery after spinal cord injury depends on whether reorganizing descending circuits form adaptive, task-specific relays rather than maladaptive ones. The corticospinal tract undergoes competitive, activity-dependent refinement during a defined juvenile window, so the timing and polarity of cortical activity may dictate whether injured circuits rewire usefully. Whether biasing cortical activity during this window can steer reorganization toward recovery, and whether naturalistic rehabilitation supplies the activity such steering requires, remain unknown.

In juvenile rats we delivered excitatory or inhibitory chemogenetic receptors, or a control vector, to the forelimb motor cortex at postnatal day 5, induced a right cervical hemisection at postnatal day 14, and gave oral clozapine-N-oxide during nightly rough-and-tumble play from postnatal day 20 to 50. Play, the rehabilitation common to all groups, did not differ between them, isolating cortical-activity polarity as the manipulated variable. Outcomes spanned forelimb dexterity, reaching kinematics, cortical output by intracortical microstimulation, descending architecture by intersectional viral tracing, and frequency-dependent depression of the Hoffmann reflex.

In untreated injured animals, cortical output re-lateralized maladaptively toward the intact forelimb, driving the ipsilateral (intact) limb in 11 of 15 evoked responses and 3 of 5 animals. Excitation preserved selective drive to the impaired forelimb (38 of 39 responses; concordant-site proportion Kruskal–Wallis H = 9.08, exact P = 0.0031), and pure ipsilateral re-lateralization was absent from all nine treated animals (3 of 5 control versus 0 of 9 treated; Fisher’s exact P = 0.027). Across behavioural, kinematic, and anatomical measures a consistent excitatory > control > inhibitory gradient emerged. Excitation enhanced wrist elevation and forward extension during reaching, restored frequency-dependent depression of the Hoffmann reflex, and selectively recruited three parallel lesion-bypassing relays — interhemispheric cortico-cortical-spinal, cortico-reticulo-spinal, and cortico-propriospinal (ipsilesional cortical relay neurons F(2,14) = 56.96, P < 0.0001). Inhibition yielded the weakest behavioural recovery and least relay recruitment, and did not restore contralesional cortical drive.

Cortical-activity polarity during a juvenile window thus acts as a developmental switch governing whether descending reorganization is adaptive or maladaptive. Guided excitation, paired with behaviourally meaningful activity, reshapes diffuse sprouting into structured, lesion-bypassing relays and corrects the spontaneous re-lateralization that otherwise misdirects recovery. These findings provide a translational rationale for activity-paired neuromodulation in paediatric spinal cord injury.

## Introduction

Traumatic spinal cord injury (SCI) severs descending commands from motor cortex to spinal circuits, producing paralysis and long-term disability. Functional recovery depends on the ability of spared pathways to reorganize and form compensatory connections. In adults, this often involves corticospinal axon sprouting rostral to the lesion to form detour circuits via propriospinal or reticulospinal relays.^1–3^ Such plasticity, however, is not uniformly adaptive. Evidence from experimental and clinical studies suggests that beneficial outcomes depend on the reorganizing circuits respecting the modular logic of motor control, relaying concerted motor commands to appropriate motor neuron pools.^4–7^ Diffuse recruitment can result in maladaptive synergies leading to hyperreflexia, spasticity, and clonus rather than task-specific movement.^8^

The corticospinal tract (CST) is uniquely sensitive to activity during early development. In rodents, CST axons transiently innervate both sides of the spinal cord before pruning to mature crossed projections during the second postnatal week.^9,10^ Perturbations of activity during this competitive period disrupt functionality, with unilateral lesions near postnatal day 14 (P14) inducing widespread contralesional CST sprouting into inappropriate spinal laminae, while lesions at later ages produce less aberrant reorganization and fewer deficits.^11^ This heightened plasticity reflects the intrinsically elevated growth status of P14 CST neurons, an altered inflammatory response, and a relative absence of myelin-associated inhibitors such as Nogo-A, MAG, and OMgp.^12–14^ These findings underscore that the timing of injury relative to CST maturation, and the activity present during this critical window, determine whether plasticity is adaptive or maladaptive.

Neuromodulation studies have defined how patterned stimulation restores function after SCI: epidural electrical stimulation is thought to activate proprioceptive afferents and re-engage latent sensorimotor circuits, strengthening Ia afferent–motoneuron synapses, recruiting propriospinal and brainstem relays that re-route commands around the lesion, and integrating cortical drive with proprioceptive feedback.^15,16^ In juvenile rodents, rough-and-tumble play provides naturalistic, patterned sensorimotor activity that engages bilateral cortical and spinal circuits, making it a compelling but mechanistically underexplored rehabilitative modality.^17–19^ We explored whether directly modulating cortical activity during the juvenile plasticity window can strengthen connectivity and facilitate adaptive outcomes.

Epidural stimulation combined with rehabilitation amplifies residual CST drive and improves functional gains,^20,21^ and recent work shows that spinal stimulation entrains motoneuron firing while supraspinal input determines whether pulses become action potentials.^22^ Emerging non-invasive modalities such as transcranial focused ultrasound add spatially precise, bidirectional modulation of deep circuits.^23^ These approaches converge on a common principle: appropriately patterned activity can steer reorganization toward useful function.

At the molecular level, adult CNS axon growth is constrained by intrinsic brakes (e.g., PTEN–mTOR, CaVα2δ) and extrinsic inhibitors.^16^ Releasing these brakes rekindles growth, but functional benefit typically requires concurrent activity to steer nascent connections toward appropriate targets:^24,25^ regenerating neurons across complete lesions restored walking only when guided to natural targets,^26^ and gabapentinoid treatment after stroke enhanced CST sprouting and forelimb recovery while suppressing maladaptive excitability.^27^ Cell-based strategies are advancing in parallel.^28^

Chemogenetics provides a uniquely selective tool to test causal, temporally precise excitation or inhibition of defined cortical populations.^29^ DREADDs are engineered G-protein– coupled receptors activated by otherwise inert ligands, giving inducible, bidirectional control of excitability with spatial and temporal fidelity — ideal for probing how the level and polarity of cortical activity shape recovery. Chemogenetic activation can directly enhance axonal growth: excitatory stimulation of dorsal root ganglion neurons promoted sensory regeneration,^30^ chemogenetic activation of CST neurons with deletion of growth-inhibitory pathways enhanced forelimb recovery after dorsal column injury,^31^ and excitatory stimulation after thoracic contusion improved locomotion in adult rats.^32^ Whether cortical excitability can be bidirectionally modulated during the juvenile window to steer descending reorganization after cervical injury remains unexplored.

Here, we used a juvenile rat model of cervical SCI aligned with the CST’s competitive period. We delivered excitatory or inhibitory DREADDs (or reporter control) to motor cortex at P5, performed right-sided C5 hemisections at P14, and actuated DREADDs during nightly play from P20–P50. Ten days after washout, we assessed recovery across behaviour, kinematics, cortical mapping, spinal reflex physiology, and histology. Across all domains an excitatory > control > inhibitory ordering emerged, separating excitation robustly from both groups while the control-versus-inhibitory difference was assay-dependent: excitatory stimulation expanded contralesional forelimb maps, restored frequency-dependent depression, and enhanced reaching, with anatomical evidence of CST sprouting onto reticulospinal and propriospinal relays, whereas inhibition markedly reduced cortical responsiveness and worsened outcomes.

By coupling excitation to rehabilitative play, we found that descending reorganization can be guided to form adaptive relays and promote functional recovery. This framework highlights a translational application: targeted augmentation of cortical output could direct CST remodelling in paediatric SCI and inform early-life neuromodulation strategies.

## Materials and methods

### Animals and experimental design

All experiments were approved by the Institutional Animal Care and Use Committee (IACUC) at Temple University (protocol# 5098) and conformed to NIH guidelines for the care and use of laboratory animals. The study is reported in accordance with the ARRIVE 2.0 guidelines. Male and female Sprague–Dawley rats (Charles River) were used across P5–P60. Because anogenital sexing at P5 adds stress incompatible with this neonatal protocol,^33^ animals were randomized without sex pre-stratification (post-hoc: 55% male, 45% female); the study was not powered for sex effects, which future work should address.

Pups were randomly assigned at P5 to one of three groups: Excitatory (AAV2-hSyn-hM3Dq-mCherry), Inhibitory (AAV2-hSyn-hM4Di-mCherry), or Control (AAV2-hSyn-mCherry). A separate cohort of naïve, uninjured animals (Normal) served as a reference for kinematic analyses. Sample sizes were determined by a priori power analysis (α = 0.05, 1–β = 0.80, effect size f = 0.80 based on IBB data), yielding a minimum of six animals per group. Final group sizes per assay were as follows. The animal was the unit of analysis throughout, except for the reaching kinematics, where the individual reach was the unit of analysis: IBB n = 18 Excitatory, 10 Control, 10 Inhibitory; Grooming and Cylinder Touch n = 13 Excitatory, 10 Control, 10 Inhibitory; Horizontal Ladder n = 6 Excitatory, 6 Control, 8 Inhibitory. H-reflex n = 8 Excitatory, 9 Control; ICMS mapping n = 4–5 per group; Intersectional tracing sample sizes across anatomical levels: cortex n = 6 Excitatory / 5 Control / 6 Inhibitory; medulla n = 8 / 7 / 8; C3/C4 spine n = 9 / 7 / 8. Play cohorts (Fig. 2A–C) were independent of those above: n = 8 Normal (uninjured) and n = 6 Lesioned animals, randomly paired. N-vs-L comparisons comprised 69 bouts and N-vs-N comparisons 42 bouts; per-animal bout counts and mean scores are provided in Source Data. Housing-manipulation cohorts (Fig. 2D–E), independent of the Fig. 3 excitatory animals, comprised n = 8 group-housed and n = 10 single-housed injured excitatory hM3Dq animals scored at the animal level.

Animals were housed under controlled conditions (12 h light/dark cycle, 22 ± 2 °C, 45– 65% humidity) with ad libitum access to standard chow and water. No animals were excluded from analysis except where post-hoc histology confirmed incomplete hemisection (residual ipsilesional PKCγ immunoreactivity at the lesion epicentre) or cortical injection misplacement (absence of reporter fluorescence in motor cortex layer V).

Post-operative care and monitoring. Following survival surgery, animals received post-operative analgesia (carprofen), antibiotic prophylaxis (cefazolin), and subcutaneous warmed saline for hydration. Animals were monitored daily for body weight, wound integrity, hydration, posture, and spontaneous behaviour, and were assessed against pre-specified humane endpoints (sustained weight loss, unrelieved pain or distress, autophagia of the affected forelimb, or failure to thrive); animals meeting these criteria were removed from the study and humanely euthanized.

### Viral vector production and validation

AAV vectors were produced in-house using a helper-virus–free triple-transfection protocol. Constructs included AAV2-hSyn-hM3Dq-mCherry, AAV2-hSyn-hM4Di-mCherry, AAV2-hSyn-mCherry (control reporter), AAV1-hSyn-nucCreR-eGFP (anterograde transsynaptic tracer), retrograde AAV2-CAG-mTagBFP (Cre-independent retrograde reporter), and retrograde AAV2-EF1α-DIO-mCherry (Cre-dependent retrograde reporter).^34^

HEK293T cells were transfected with the transgene, packaging and helper plasmids, and virus was purified by double caesium-chloride ultracentrifugation and dialysis as previously described.^14,35^ AAV2-eGFP used in tracing controls was produced via the Bodhi CGT platform (Appraiseye Inc., Philadelphia, PA).

Genomic titres were determined by qPCR against AAV2 ITR standards (all vectors >1 × 10¹3 genome copies mL⁻¹), and reporter expression was confirmed in vitro before use.

### Neonatal cortical injections

At P5, pups were anaesthetized with hypothermia-assisted ketamine (40 mg kg⁻¹, i.p.) and secured in a neonatal stereotaxic frame. Viral suspension was delivered to the left forelimb motor cortex at three sites following our published surgical protocol (coordinates and volumes in the Supplementary material).^36^ The pipette was retained in situ for 3 min post-injection to minimize reflux. Pups were returned to the dam on a warming pad and monitored until ambulatory.

### Cervical spinal cord hemisection

At P14, pups were anaesthetized with ketamine/xylazine (90/10 mg kg⁻¹, i.p.) and secured in a custom spinal stabilization frame. A midline dorsal incision exposed the cervical vertebrae, and a right-sided C5 laminectomy was performed under 20× magnification. A complete right hemisection was performed using fine iridectomy scissors, extending the incision to the midline until the central canal and contralateral dorsal column were clearly visualized, ensuring complete interruption of the right corticospinal tract and all ipsilateral descending pathways.^37^ Musculature was closed in layers and skin with wound clips. Lesion completeness was verified post-hoc by PKCγ immunolabelling of corticospinal tract axons at thoracic levels (∼T4/T5); animals showing any residual ipsilesional PKCγ-positive fibres in the dorsal funiculus were excluded.

### Rehabilitative play paradigm and CNO delivery

From P20 to P50, animals were housed in groups of three same-sex littermates to allow natural rough-and-tumble play, a behaviour that engages bilateral cortical and subcortical motor networks. Night-time play sessions were video recorded under red-light illumination for 20 min weekly. Play behaviour was quantified using an ethogram modified from Whishaw et al.^38^ that classifies events including dorsal contacts, nape attacks, pinning, and forelimb boxing.^39^ Recordings were scored by two blinded observers. To isolate the rehabilitative contribution of social play, an additional cohort of injured excitatory animals was housed individually or in groups under otherwise identical conditions and CNO supplementation.

Chemogenetic activation was achieved by supplementing DietGel Recovery (ClearH₂O, Lancaster, PA) with clozapine-N-oxide (CNO; Tocris, Cat# 4936; 3 mg kg⁻¹ day⁻¹) from P20 to P50.^40,41^ All three groups (Excitatory, Inhibitory, Control) received identical CNO-supplemented gel to control for any off-target effects of the ligand. The hM3Dq and hM4Di constructs are extensively validated, and oral CNO produces sustained brain bioavailability sufficient for DREADD activity in rodents.^29,40,41^ Animals were drug-free for ≥10 days before behavioural testing to ensure that all assessments reflected stabilized plasticity rather than acute pharmacological effects.

### Behavioural assays

Behavioural assessment was performed at P60, ≥10 days after the final CNO dose, by experimenters blinded to group allocation. Four assays were conducted in a fixed order across two consecutive days.

IBB forelimb recovery scale: Forelimb dexterity was scored on the 0–9 Irvine–Beattie– Bresnahan (IBB) scale from video of cereal manipulation, independently for each limb.^42^

Grooming assay: Forelimb range of motion was scored on a 0–5 scale using a grooming task, recording the highest landmark reached by the affected forepaw.^43^

Cylinder rearing test: Spontaneous affected-forepaw use was scored over the first 20 weight-supporting rears in a glass cylinder, using the cylinder test.^44^

Horizontal Ladder: Forelimb placement was assessed on a horizontal ladder (1 m, 10 mm rung spacing) over three traversals, expressed as percentage correct placements.^45^

### Kinematic analysis of skilled reaching

Rats were trained to retrieve food pellets through a 10 mm vertical slit, and reaches were recorded on high-speed video (250 frames s⁻¹); acquisition details are given in the Supplementary material.^46^

Shoulder, elbow, wrist and digit-tip markers were tracked with DeepLabCut (network details in the Supplementary material).^47^ Coordinates were referenced to the slit plane (+x forward, +y vertical), and each reach was temporally normalized and aligned to the frame of peak forward extension.

Kinematic parameters included maximum reach distance (Δx from shoulder), maximum wrist elevation (Δy), and wrist–shoulder separation in both axes. All trajectories underwent quality control for occlusions and low-confidence frames. Per-group n_reaches were 13 / 11 / 7 / 4 (Excitatory / Control / Inhibitory / Normal) for wrist-coordinate endpoints (Fig. 4); analyses requiring shoulder or elbow markers (joint extrema, Fig. S1; wrist–shoulder separation, Fig. S2) excluded one Control reach lacking trackable shoulder/elbow coordinates, giving 13 / 10 / 7 / 4. Between-group differences in time-varying kinematics (Δx(t) and Δy(t) wrist–shoulder trajectories) were evaluated using one-dimensional statistical parametric mapping (SPM1d)^48^ with two-sample t-tests and random-field–theory correction (α = 0.05, two-tailed). Scalar kinematic endpoints were compared using Kruskal–Wallis tests with Holm-corrected post-hoc Mann–Whitney U tests.

### Intracortical microstimulation mapping

At the study endpoint (P60), animals were anaesthetized with ketamine (100 mg kg⁻¹, i.p.) supplemented as needed to maintain areflexia and positioned in a stereotaxic frame. A bilateral frontal craniotomy exposed the agranular motor cortices (AP +4.0 to −2.0 mm from bregma; ML 0.5–4.0 mm). The skull was thinned with a high-speed drill under saline irrigation, the dura was removed under microscopic visualization, and the cortical surface was kept moist with warm saline throughout.

At each penetration (layer V; electrode and stimulation parameters in the Supplementary material),^49,50^ the minimum current evoking a visible movement in ≥50% of trains (25 µA steps, 50–300 µA) defined the movement threshold; sites unresponsive at 300 µA were classified as non-responsive.

Sites were mapped on a 0.5 × 0.5 mm bilateral grid, and evoked movements were categorized (forelimb, vibrissae, facial, trunk, or hindlimb) by two blinded observers. For each animal, the proportion of sites evoking forelimb responses was calculated separately for the contralesional (left) and ipsilesional (right) cortex. For forelimb responses evoked from the contralesional cortex, the side of the evoked movement was scored and each response classified as contralateral (right, impaired forelimb), ipsilateral (left, intact forelimb), or bilateral; recruitment of the ipsilesional cortex into right-forelimb responses was scored per animal. To assess the contribution of interhemispheric pathways, muscimol (0.5 µL, 5 mM; Tocris) was injected into the ipsilesional cortex while contralesional stimulation continued, and any abolition of evoked responses was documented.

### H-reflex electrophysiology

H-reflexes were recorded bilaterally from the forelimbs of anaesthetized rats (ketamine, 90 mg kg⁻¹ i.p.) immediately prior to ICMS mapping, stimulating the ulnar nerve at the wrist and recording from the abductor digiti minimi, as previously described.^51^

The H-reflex threshold was defined as the current evoking a stable response in ≥50% of trials, and recruitment curves were acquired to 2× threshold.

Frequency-dependent depression (FDD) was assessed by delivering paired stimuli at interstimulus intervals ranging logarithmically from 10 s (0.1 Hz) to 0.01 s (100 Hz).^51,52^ For each frequency, 25 paired trials were collected and averaged. The area under the curve (AUC) of the test H-wave was normalized to the conditioning H-wave AUC to compute the degree of reflex suppression. FDD served as a surrogate for presynaptic inhibition: reduced suppression indicated pathological hyperreflexia, and increased suppression indicated restoration of GABAergic inhibitory tone.^53,54^ Inhibitory (hM4Di) animals also underwent H-reflex recording, but these data were acquired on a separate acquisition system and were not frequency-matched to the excitatory and control recordings; they were therefore excluded from quantitative FDD analysis, and the reported H-reflex cohort comprises excitatory and control animals only.

### Intersectional viral tracing

Following completion of all behavioural, kinematic, and electrophysiological assessments, animals underwent dual tracing surgery to map cortico–subcortical–spinal relay circuits. Under ketamine/xylazine anaesthesia (90/10 mg kg⁻¹ i.p.), a craniotomy over the left (contralesional) forelimb motor cortex was used to deliver AAV1-hSyn-nucCreR-eGFP across four sites (coordinates and volumes in the Supplementary material).^14^

For retrograde labelling, a laminectomy exposed the right (ipsilesional) C6–T1 cord, and a 1:1 mixture of retrograde AAV2-mTagBFP (Cre-independent) and AAV2-DIO-mCherry (Cre-dependent) was injected into the intermediate grey matter at ten sites along the rostro– caudal axis (coordinates in the Supplementary material).

In this intersectional design, BFP labels all neurons with projections reaching the caudal spinal cord regardless of cortical input, while mCherry expression is restricted to the subset that received transsynaptic Cre from cortical neurons. The proportion of mCherry⁺ (red-labelled) neurons among total BFP⁺ retrograde labelling, provides a readout of cortical relay engagement. Quantification was performed in three regions: ipsilesional primary motor cortex, contralesional medullary reticular formation (gigantocellularis [Gi] and lateral paragigantocellularis [LPGi] nuclei), and ipsilesional C3/C4 propriospinal interneurons.

### Histology and immunohistochemistry

At P60, animals were euthanized with sodium pentobarbital (100 mg kg⁻¹, i.p.) and transcardially perfused with 4% paraformaldehyde, and brains and spinal cords were cryosectioned at 30 µm as previously described.^14^

Native fluorescence of mCherry, GFP, and BFP was visualized directly in mounted sections. For lesion verification and CST integrity, free-floating sections were immunostained with anti-PKCγ (1:200; Abcam, Cat# ab317315, clone EPR28643-68, RRID:AB_3720317) and anti-GFAP (1:1000; Dako, Cat# Z0334, RRID:AB_10013382), followed by species-appropriate Alexa Fluor–conjugated secondary antibodies (1:1000; Invitrogen). Sections were mounted on SuperFrost Plus slides with Fluoromount-G (SouthernBiotech) and imaged on a Zeiss Axio Imager.M2 with ApoTome.2 structured illumination, analysed using ZEN software (Carl Zeiss). Lesion completeness was confirmed by absence of PKCγ-positive fibres in the ipsilesional dorsal funiculus at thoracic levels.

For motor cortex and medullary sections, discrete cell bodies positive for mCherry (red), mTagBFP (blue), or eGFP (green) were counted manually within standardized regions of interest. Cortical ROIs encompassed layers V–VI of the agranular forelimb motor area, defined by cytoarchitectural landmarks. Medullary ROIs encompassed the Gi and LPGi nuclei, delineated according to the Paxinos and Watson rat brain atlas.^55^ For each animal, counts were summed across three to four sections per animal per region sampled at 200 µm rostro-caudal intervals. The percentage of mCherry+ neurons among total BFP+ retrogradely labelled neurons was calculated per animal as the primary readout of cortical relay engagement.

At the C3/C4 spinal cord rostral to the lesion, quantification was performed on fluorescent label area.^1,2,56,57^ mCherry and BFP channels were independently thresholded at a fixed intensity cutoff (120–255, 8-bit) applied across all images and animals. The spinal grey matter ROI was manually outlined in FIJI using the freehand selection tool,^58^ and the resulting binary masks were measured within this ROI. Total R+B area was calculated as the union of mCherry-positive and BFP-positive pixels; mCherry+ area was expressed as a percentage of this total per animal.

### Statistical analysis

Data were analysed using GraphPad Prism 10 (GraphPad Software, San Diego, CA), MATLAB R2023b with the SPM1d toolbox (www.spm1d.org), R v4.2.3, and Python (spm1d, scipy.stats, statsmodels). Group values are reported as mean ± SD in the Results text and Supplementary Tables; figure error bars represent mean ± SEM unless otherwise noted, with individual data points overlaid. Normality and homogeneity of variance were assessed within each group using the Shapiro–Wilk and Brown–Forsythe tests, respectively. Test selection was data-driven: one-way ANOVA with Tukey’s post-hoc when both assumptions held, Kruskal– Wallis with Dunn’s post-hoc when normality was violated, and Welch’s ANOVA with Games– Howell post-hoc when variances were unequal.

Behavioural scores were averaged within animal across two test sessions, with the animal as the unit of analysis. The horizontal ladder was added to the testing battery after the excitatory and control cohorts had completed the other three assays, accounting for the smaller n in those groups. IBB, Grooming, and Cylinder Touch (non-normal distributions) were analysed with Kruskal–Wallis and Dunn’s post-hoc; Horizontal Ladder (both assumptions met) with one-way ANOVA and Tukey’s HSD. Play two-group comparisons used two-tailed unpaired t-tests, except for the Normal vs. Lesioned play score comparison, where variance heterogeneity (F-test p < 0.0001) prompted Welch’s correction. For Normal vs. Lesioned and Normal vs. Normal play comparisons, the unit of analysis was the individual play bout, reflecting the dyadic structure of rough-and-tumble play in which each interaction yields an independent observation of pin attempts and engagement.^38,39^

Intersectional tracing proportions used one-way ANOVA with Tukey’s correction (cortex and medulla, equal variances) or Welch’s ANOVA with Games–Howell correction (C3/C4, heterogeneous variance); total neuron counts used one-way ANOVA or Welch’s ANOVA with Games–Howell post-hoc where Brown–Forsythe indicated heterogeneous variance; ICMS responsiveness used Kruskal–Wallis with Dunn’s correction; categorical comparisons of response laterality and ipsilesional-cortex recruitment used Fisher’s exact tests. Kinematic scalar endpoints used Kruskal–Wallis with Holm-corrected post-hoc Mann– Whitney U tests; time-varying kinematics used SPM1d two-sample t-tests with random-field– theory correction. Per-cluster SPM statistics are in Tables S1, S4, S5; extremal-measure statistics in Tables S2, S3; Fig. 6 cell counts in Fig. S6.

Terminal intracortical microstimulation mapping was completed in a subset of animals (n = 4–5 per group), below the a priori target of six, reflecting attrition across the extended juvenile protocol and the technical demands of terminal mapping. The microstimulation comparisons are therefore the most conservatively powered in the study. To limit the influence of small per-group samples, cortical output was analysed at the level of the animal — the per-animal proportion of concordant contralesional sites — rather than by pooling individual stimulation sites, avoiding pseudoreplication, and group differences were evaluated with exact non-parametric tests.

For H-reflex, two-way repeated-measures ANOVA was used for bilateral within-group comparisons (limb × frequency) and two-way mixed ANOVA for between-group comparisons within a single limb (group × frequency). Greenhouse–Geisser correction was applied to within-subject factors where sphericity assumptions were not met; significant interactions were followed by Bonferroni-corrected post-hoc comparisons at each frequency. Effect sizes are reported as ηZG (within-subject ANOVA), ηZp (mixed ANOVA), or Cohen’s d (pairwise). H-reflex analyses were performed in Python using pingouin v0.5 and SciPy.

Kinematic trajectory pre-processing (interpolation, temporal alignment and resampling, calibration to millimetres, and low-pass filtering of inhibitory elbow trajectories) is detailed in the Supplementary material; significant SPM clusters were reported as contiguous temporal intervals.

Statistical significance was set at α = 0.05. All data collection and statistical analyses were performed by investigators blinded to group allocation.

The study was exploratory and was not preregistered; no analysis plan was lodged in a public registry before data collection.

## Results

Excitatory (hM3Dq), inhibitory (hM4Di) or control (mCherry) DREADDs were expressed in the left forelimb motor cortex at P5, followed by a right C5 hemisection at P14 that induces forelimb deficits persisting into adulthood, and oral CNO throughout the juvenile play window (P20–P50; Fig. 1). Group-housed juvenile rats were used because rough-and-tumble play is instinctual, robust, and generates quantifiable complex forelimb movements that increase locomotor recovery after SCI.^7,17,18,38,59,60^

**Figure 1.**
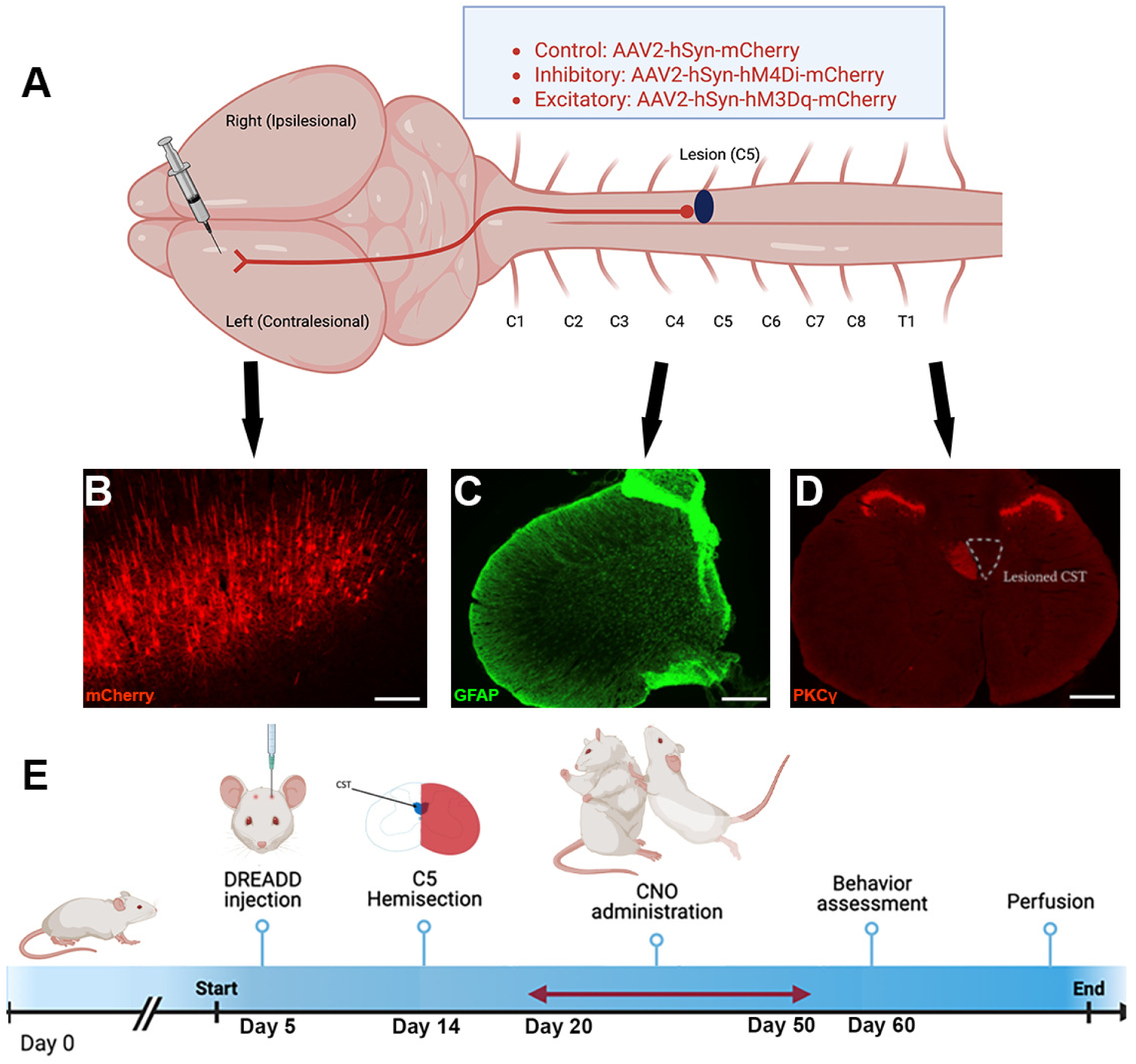
Experimental design and verification of cortical injection and lesion completeness. **(A)** Schematic of viral injection into the left forelimb motor cortex at P5. Excitatory (AAV2-hSyn-hM3Dq-mCherry), inhibitory (AAV2-hSyn-hM4Di-mCherry), or control (AAV2-hSyn-mCherry) constructs were delivered across three sites (AP +0.5 mm; ML −1.0, −1.5, −2.0 mm; depth −0.6 mm from cortical surface). **(B)** Representative coronal section showing mCherry expression spanning forelimb motor cortex layers V–VI; scale bar 200 µm. **(C)** GFAP immunolabelling (green) at the C5 hemisection epicentre, confirming complete right-sided cord interruption to the midline; scale bar 500 µm. **(D)** PKCγ immunolabelling at T4/T5 confirming absence of corticospinal axons in the right (ipsilesional) dorsal funiculus (dotted outline indicates the lesioned right corticospinal tract); animals with residual ipsilesional PKCγ-positive fibres were excluded. Scale bar 500 µm. **(E)** Experimental timeline: P5 cortical AAV injection; P14 right C5 hemisection; P20–P50 nightly oral CNO via DietGel during rough-and-tumble play; behavioural, kinematic, ICMS, H-reflex, and intersectional tracing assessments at P60, ten days after CNO washout. Days on the timeline correspond to postnatal age (e.g., Day 5 = P5). Alt text: Diagram of the study design and timeline — P5 cortical AAV injection, P14 right C5 spinal hemisection, oral CNO during nightly play from P20 to P50, and assessments at P60 — shown with fluorescence micrographs confirming mCherry expression in motor cortex layers V–VI, GFAP labelling at the C5 lesion, and absence of PKCγ-positive corticospinal axons in the right dorsal funiculus.

### Cervical SCI impairs play competence while sparing engagement

Lesioned rats frequently attempted nape attacks with the affected limb but were unable to complete pins, consistent with impaired forelimb control. Across paired bouts they scored significantly fewer pin victories than normal rats (Normal 1.35 ± 0.16 vs. Lesioned 0.77 ± 0.09 per bout; Fig. 2A; Supplementary Video 1), whereas bouts between normal cage-mates were balanced (Fig. 2B), and lesioned rats trended toward elevated overall activity (Fig. 2C). Chemogenetic group assignment did not alter play engagement among injured cohorts, excluding altered play as a confound of behavioural recovery (Fig. S7).

**Figure 2.**
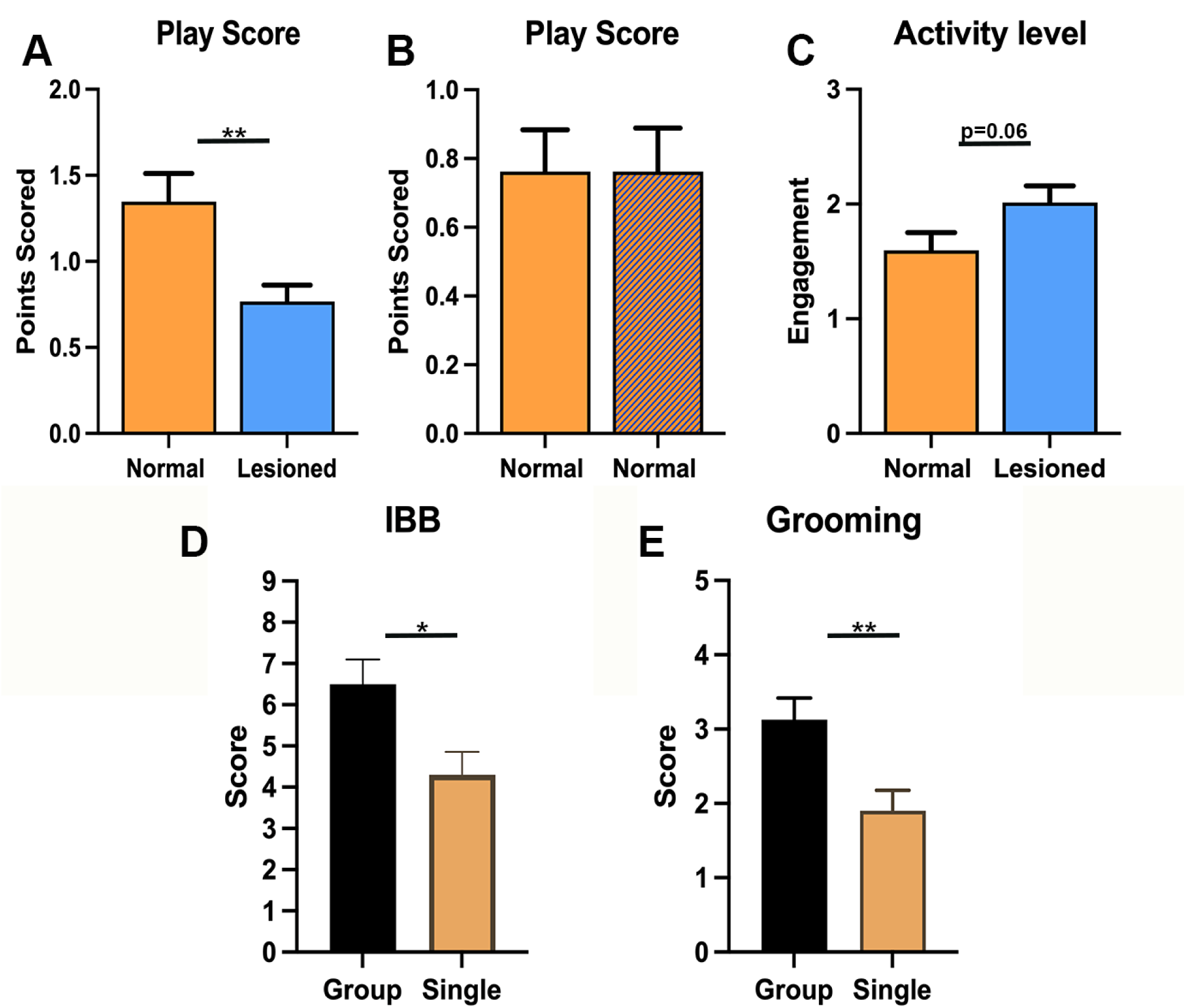
Rough-and-tumble play behaviour during the juvenile actuation window. **(A)** Pin victories scored during inter-group bouts between lesioned and normal rats; lesioned rats achieved fewer pins, consistent with impaired forelimb control (Normal 1.35 ± 0.16 vs. Lesioned 0.77 ± 0.09 per bout; t(109.6) = 3.08, p = 0.003, Cohen’s d = 0.52). **(B)** Among normal rats paired with normal cage-mates, bouts were balanced with no individual achieving higher pin counts (t(82) = 0.000, p > 0.99). **(C)** Lesioned rats showed elevated overall activity (combined social and non-social exploration with play engagement) relative to normal rats (Normal 1.60 ± 0.16 vs. Lesioned 2.01 ± 0.14; t(109) = 1.90, p = 0.061). **(D, E)** Forelimb recovery in injured excitatory DREADD animals housed in groups of three for nightly rough-and-tumble play (group-housed) versus injured excitatory DREADD animals housed singly (single-housed) throughout the P20–P50 actuation window, assessed by IBB cereal-manipulation (D; Group 6.50 ± 0.60 vs. Single 4.30 ± 0.56; t(16) = 2.68, p = 0.017, Cohen’s d = 1.27) and grooming reach (E; Group 3.13 ± 0.30 vs. Single 1.90 ± 0.28; t(16) = 3.01, p = 0.008, Cohen’s d = 1.43). Group-housed animals showed significantly greater recovery on both assays, demonstrating that play participation is required for the full behavioural benefit of excitatory chemogenetic stimulation. Bars represent mean ± SEM with individual data points overlaid. Two-tailed unpaired t-tests for B–E; Welch’s t-test with Satterthwaite-corrected df for A (variance heterogeneity, F-test p < 0.0001). ** p < 0.01, *** p < 0.001. Alt text: Bar graphs of rough-and-tumble play showing that lesioned rats score fewer pin victories than normal rats and trend toward higher overall activity, that normal-versus-normal bouts are balanced, and that group-housed excitatory animals outperform singly-housed ones on the IBB and grooming scores, with individual data points overlaid.

To test whether play, rather than excitation alone, drove recovery, injured excitatory animals were housed individually or in groups throughout the actuation window. Group-housed excitatory animals significantly outperformed their singly-housed counterparts on both IBB (Fig. 2D) and grooming (Fig. 2E), with large effect sizes for both (Cohen’s d > 1.2). Excitatory chemogenetic stimulation therefore required co-occurring social play to confer its full behavioural benefit.

### An excitatory > control > inhibitory gradient across forelimb behavioural recovery

Ten days after the final CNO dose, forelimb function was tested off-drug, so that changes reflected stabilized plasticity rather than acute activity (Fig. 3). Hand and wrist dexterity improved markedly with excitatory DREADDs on the IBB scale (Kruskal–Wallis p < 0.0001; Fig. 3A),^42^ whereas inhibitory animals manipulated the treat strictly with the unaffected forelimb. The same excitatory > control > inhibitory hierarchy was seen for grooming reach (p < 0.0001; Fig. 3B),^43^ spontaneous affected-forepaw use in the cylinder (p < 0.0001; Fig. 3C), and forelimb placement on the horizontal ladder (one-way ANOVA p = 0.0008; Fig. 3D). Control and inhibitory animals did not differ on the cylinder or ladder (Dunn’s p > 0.99 and Tukey’s p = 0.96, respectively). Effect sizes for the excitatory comparisons were large throughout (Cohen’s d vs. control: IBB 1.79, grooming 1.80, cylinder 1.67, ladder 2.22; vs. inhibitory: IBB 5.09 (inflated by near-zero inhibitory variance), grooming 3.03, cylinder 2.18, ladder 2.83), and control animals significantly exceeded inhibitory animals on both graded dexterity measures (Dunn’s: IBB p = 0.048, grooming p = 0.045).

**Figure 3.**
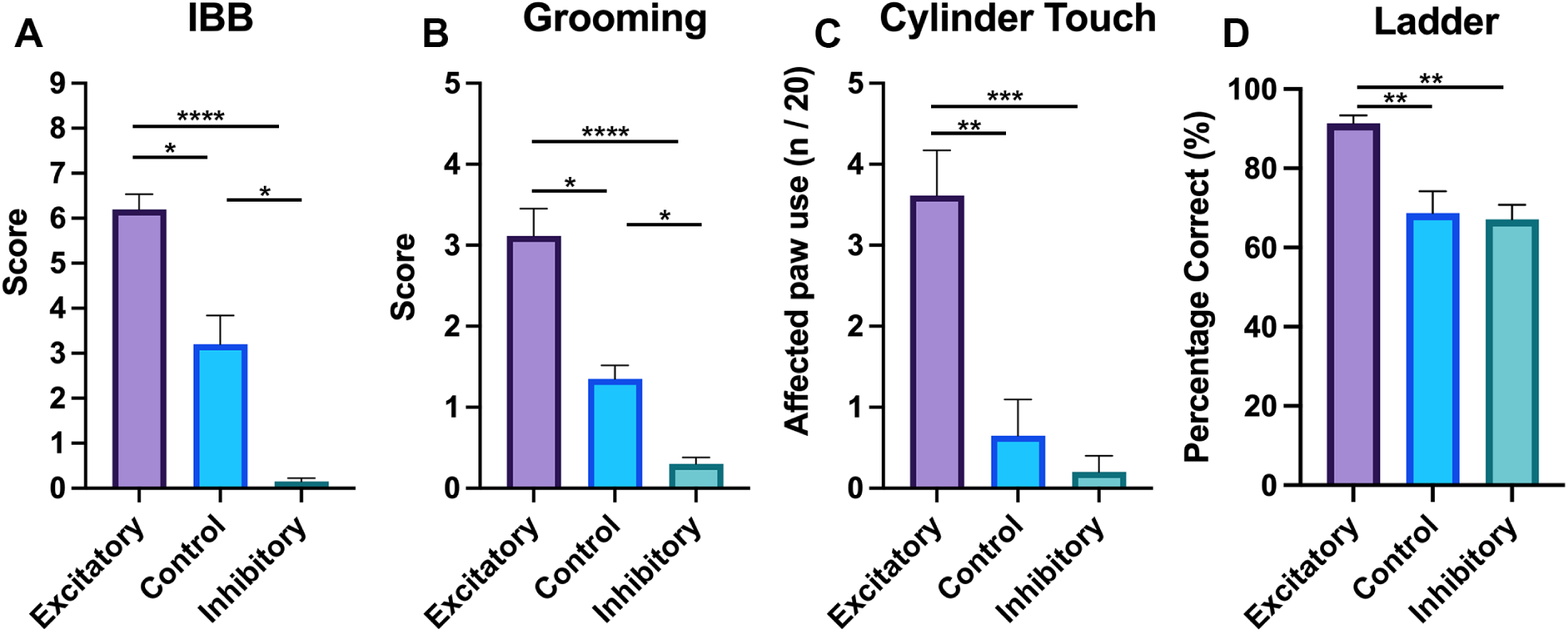
Forelimb behavioural recovery shows a consistent excitatory > control > inhibitory hierarchy. Each animal’s score is the mean of two test sessions. **(A)** IBB cereal-manipulation scores (Excitatory 6.19 ± 0.34, median 6.25 [5.00–7.63]; Control 3.20 ± 0.64, median 3.00 [1.38–4.75]; Inhibitory 0.15 ± 0.08, median 0.00 [0.00–0.50]; Kruskal–Wallis H = 27.35, p < 0.0001, εZ = 0.72; Dunn’s post-hoc: Excitatory vs. Control p = 0.042, Excitatory vs. Inhibitory p < 0.0001, Control vs. Inhibitory p = 0.048). **(B)** Grooming reach scores on a 0–5 ordinal scale (Excitatory 3.12 ± 0.34, median 3.00 [2.00–4.00]; Control 1.35 ± 0.17, median 1.25 [1.00–2.00]; Inhibitory 0.30 ± 0.08, median 0.50 [0.00–0.50]; Kruskal–Wallis H = 25.94, p < 0.0001, εZ = 0.80; Dunn’s post-hoc: Excitatory vs. Control p = 0.038, Excitatory vs. Inhibitory p < 0.0001, Control vs. Inhibitory p = 0.045). **(C)** Cylinder exploration: number of rears (out of 20) in which the affected forepaw made wall contact, either alone or together with the unaffected forepaw (Excitatory 3.62 ± 0.56, median 3.50 [2.00–4.50]; Control 0.65 ± 0.45, median 0.00 [0.00–1.00]; Inhibitory 0.20 ± 0.20, median 0.00 [0.00–0.00]; Kruskal–Wallis H = 21.42, p < 0.0001, εZ = 0.65; Dunn’s post-hoc: Excitatory vs. Control p = 0.0012, Excitatory vs. Inhibitory p < 0.0001, Control vs. Inhibitory p > 0.99). **(D)** Horizontal ladder paw placement (Excitatory 91.33 ± 2.02, median 93.00 [86.75–95.50]; Control 68.67 ± 5.54, median 70.00 [57.00–81.00]; Inhibitory 67.13 ± 3.67, median 63.25 [59.63–76.25]; one-way ANOVA F(2,17) = 11.06, p = 0.0008, ηZ = 0.57; Tukey’s post-hoc: Excitatory vs. Control p = 0.0037, Excitatory vs. Inhibitory p = 0.0012, Control vs. Inhibitory p = 0.96). Bars and group values: mean ± SEM; medians [IQR] in brackets. Excitatory = purple, Control = blue, Inhibitory = teal. Sample sizes: A, n = 18 / 10 / 10; B, n = 13 / 10 / 10; C, n = 13 / 10 / 10; D, n = 6 / 6 / 8. * p < 0.05, ** p < 0.01, *** p < 0.001, **** p < 0.0001. Alt text: Four bar graphs — IBB, grooming, cylinder rearing, and horizontal ladder — each showing an excitatory greater than control greater than inhibitory pattern of forelimb recovery, with individual data points overlaid.

### Restored kinematics during skilled reaching in excitatory animals

To resolve reaching dynamics beyond these general assays, forelimb kinematics were tracked from high-speed video using DeepLabCut (Fig. 4A–F).^47^ Control and inhibitory animals failed to reach past the plexiglass opening, consistently reaching lower and laterally and hitting the wall (Fig. 4C, E). Rats that received excitatory DREADDs showed reaching distances comparable to those from normal animals (Excitatory vs. Normal p = 0.41, Mann– Whitney U) and significantly greater forelimb extension than controls (p = 0.0004; Fig. 4D). Excitatory rats also consistently reached higher than normal rats when reaching through the opening (Excitatory vs. Normal p = 0.040, Mann–Whitney U; Fig. 4F), lifting their paw well above the pellet prior to dropping down onto it. Once these rats contacted the pellet, they rarely grasped it and instead retrieved it by dragging, consistent with residual deficits in digit individuation.

**Figure 4.**
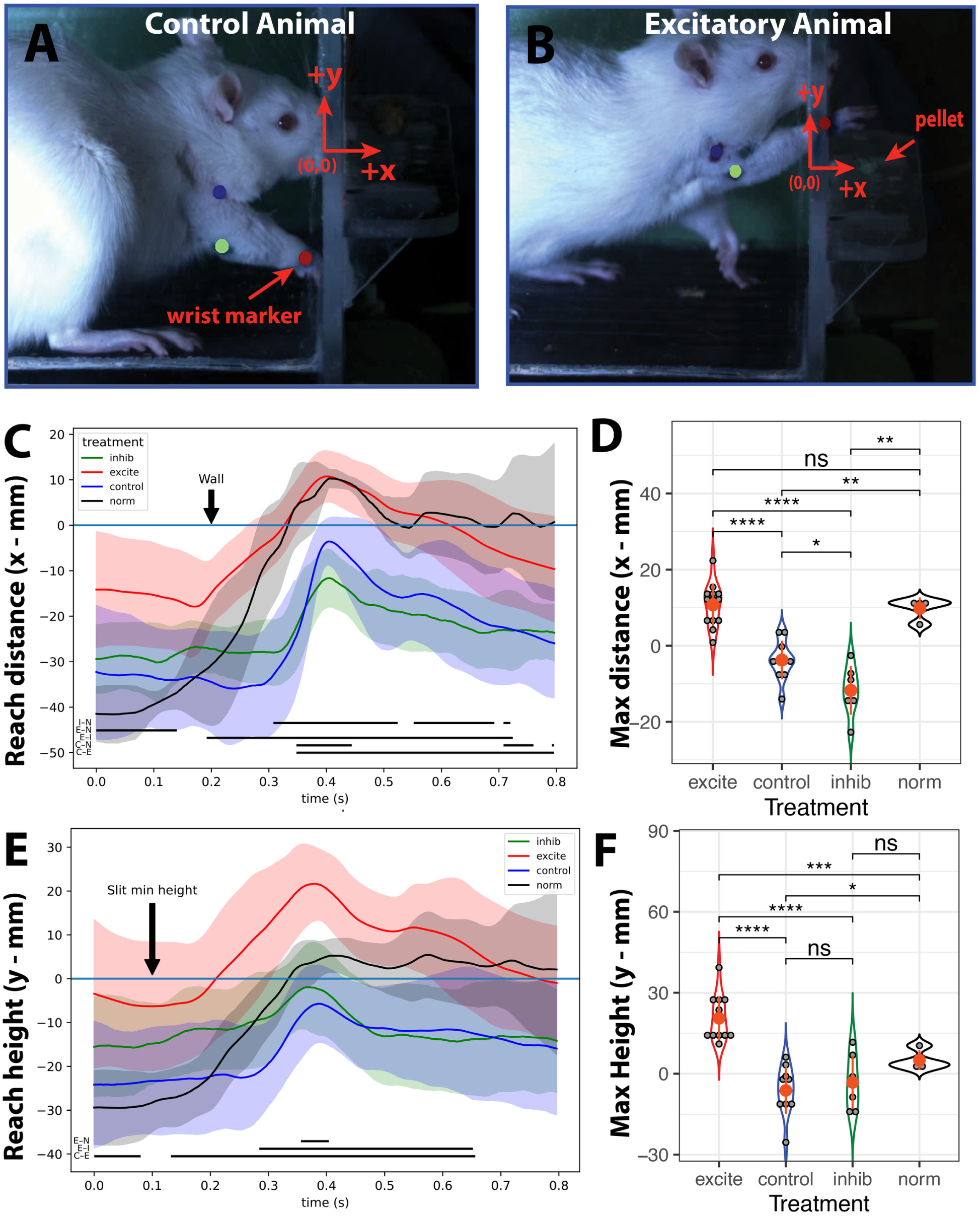
Excitatory chemogenetic stimulation enhances skilled reaching kinematics. Still images from high-speed video of representative control **(A)** and excitatory **(B)** animals reaching through a slit in a plexiglass box, with tracked anatomical markers and coordinate system overlaid (origin at slit plane; +x forward toward pellet, +y vertical elevation). To examine reach trajectory and endpoint maxima, wrist marker coordinates were extracted **(C–F)**. Wrist x- and y-coordinates against time are shown in C and E (lines = group mean; shaded ribbons = ± 1 SD). Both reach trajectories **(C, E)** and maximum coordinates **(D, F)** differed significantly across treatment groups. Black horizontal bars below each trajectory panel denote time regions of significant between-group differences (p < 0.05; one-dimensional statistical parametric mapping, two-sample t-tests with random-field-theory correction; SPM1d v0.4.0).48 Pairwise comparisons indicated by letter codes: I–N inhibitory vs. normal; E–N excitatory vs. normal; E–I excitatory vs. inhibitory; C–N control vs. normal; C–E control vs. excitatory. **(D)** Maximum wrist x (Excitatory 11.09 ± 5.41 mm; Control −2.54 ± 5.16 mm; Inhibitory −11.32 ± 6.33 mm; Normal 10.46 ± 2.08 mm; Kruskal–Wallis H = 26.51, p < 0.0001; Holm-corrected Mann–Whitney U: Excitatory vs. Control p = 0.0004, Excitatory vs. Inhibitory p = 0.0002, Excitatory vs. Normal p = 0.41, Control vs. Inhibitory p = 0.023, Control vs. Normal p = 0.006, Inhibitory vs. Normal p = 0.018). **(F)** Maximum wrist y (Excitatory 24.63 ± 8.65 mm; Control −0.54 ± 5.44 mm; Inhibitory −0.23 ± 8.68 mm; Normal 9.98 ± 7.05 mm; Kruskal–Wallis H = 25.55, p < 0.0001; Holm-corrected Mann–Whitney U: Excitatory vs. Control p = 0.0002, Excitatory vs. Inhibitory p = 0.0002, Excitatory vs. Normal p = 0.040, Control vs. Inhibitory p = 0.93, Control vs. Normal p = 0.040, Inhibitory vs. Normal p = 0.15). Colour key: Excitatory = red, Control = blue, Inhibitory = green, Normal = black. * p < 0.05, ** p < 0.01, *** p < 0.001, **** p < 0.0001. Alt text: High-speed-video stills of a control and an excitatory rat reaching through a slit with tracked joint markers, alongside wrist-trajectory and maximum-reach plots showing that excitatory animals reach farther and higher than control and inhibitory animals and comparably to uninjured animals.

Joint-coordinate and trajectory analyses (Figs S1–S3, Tables S1–S5) showed that across all injured groups the shoulder was shifted significantly toward the pellet relative to uninjured animals, indicating recruitment of the shoulder — innervated above the lesion — as a compensatory strategy, whereas uninjured animals achieved forward reach predominantly through wrist and elbow extension. Critically, excitatory animals showed significantly greater elbow horizontal extension than inhibitory animals, consistent with partial restoration of the triceps musculature innervated by C6–C8. This was corroborated by the wrist–shoulder displacement analysis: excitatory animals maintained a significantly larger ΔX and ΔY separation between wrist and shoulder throughout the reach trajectory compared with control and inhibitory animals, indicating that excitatory animals successfully extended and elevated the wrist relative to the shoulder to clear the opening.

### Excitatory stimulation expands and correctly lateralizes contralesional forelimb representation

To assess whether recovered kinematics reflected restored cortical drive to distal musculature, we performed intracortical microstimulation (ICMS) mapping of the motor cortex (Fig. 5A–C; non-forelimb responses in Fig. S4). Mapping revealed pronounced group-dependent differences in the distribution and excitability of sites evoking distal forelimb movements (Fig. 5D–F). In control rats, forelimb movements were confined to a narrow region near rostro–caudal coordinate 0 mm, consistent with the canonical forelimb motor area, and minimal responses were observed in the contralesional cortex. Inhibitory animals exhibited widespread contralesional unresponsiveness even at maximal intensities (300 µA). In contrast, excitatory animals displayed an expanded and more focally concentrated distribution of forelimb-evoked sites within the contralesional motor cortex, including robust right forelimb movements at moderate intensities (≤100 µA). Recruitment of the ipsilesional (right) cortex into right forelimb activity was observed exclusively in excitatory animals (4/5 excitatory rats vs 0/9 control and inhibitory rats; Fisher’s exact P = 0.005), consistent with partial preservation of bilateral representations and mirror-like motor coding. Injection of muscimol into the ipsilesional cortex abolished motor responses associated with this interhemispheric pathway in a representative excitatory animal. Bilateral cortical anterograde tracing in a representative excitatory animal demonstrated dense callosal projections connecting the contralesional and ipsilesional forelimb motor cortices (Fig. S8).

**Figure 5.**
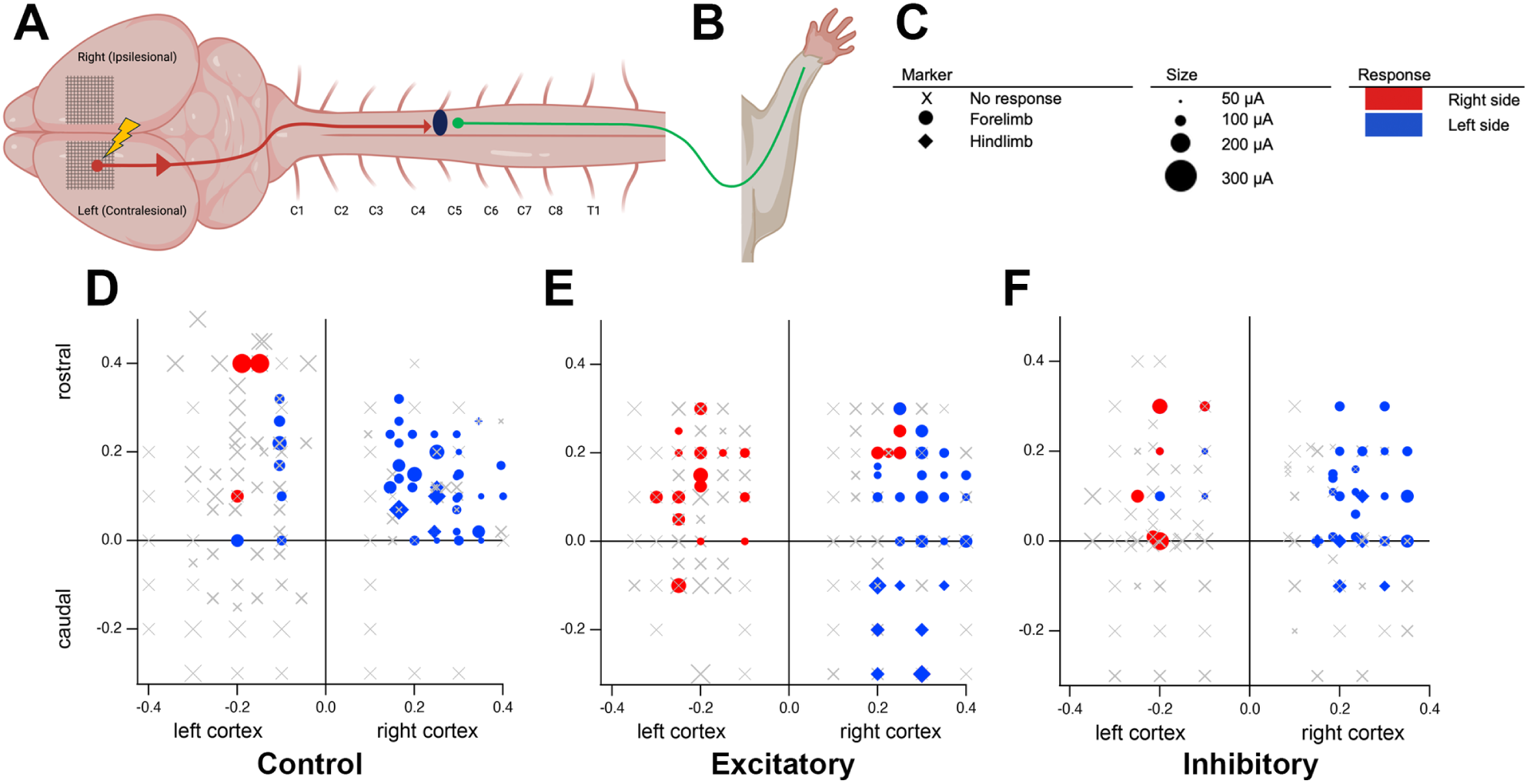
Intracortical microstimulation reveals expanded and correctly lateralized contralesional forelimb representation in excitatory animals. **(A)** Schematic showing bilateral motor cortex (left = contralesional, right = ipsilesional) and the C5 right hemicord lesion. **(B)** Forelimb illustration indicating the contralateral musculature whose evoked movement was scored at each stimulation site. **(C)** Key applying to panels D–F: marker shape encodes evoked-movement category (× = no response; ● = forelimb; ◆ = hindlimb); marker size encodes the lowest current evoking a visible movement in ≥50% of stimulation trains (50, 100, 200, 300 µA; smaller = lower threshold); marker colour encodes the side of the evoked movement (red = right side; blue = left side). **(D–F)** Representative ICMS response maps for control **(D)**, excitatory **(E)**, and inhibitory **(F)** animals, plotted with rostro–caudal coordinate on the y-axis and medio–lateral coordinate on the x-axis (left half of each panel = left cortex / contralesional; right half = right cortex / ipsilesional). Excitatory animals showed a markedly higher median proportion of concordant contralesional forelimb sites (0.21) than control (0.02) or inhibitory (0.02) animals (Kruskal–Wallis H = 9.08, exact p = 0.0031; Dunn’s post-hoc: Excitatory vs. Control p = 0.0236, Excitatory vs. Inhibitory p = 0.0367, Control vs. Inhibitory p > 0.99). Recruitment of the ipsilesional (right) cortex into right-forelimb activity was observed in 4/5 excitatory rats vs. 0/9 control and inhibitory rats (Fisher’s exact P = 0.005). Among contralesional (left) cortex forelimb responses, 11/15 in control animals drove the ipsilateral (left, intact) forelimb, in 3 of 5 control animals, indicating maladaptive re-lateralization; pure ipsilateral responses were absent from every excitatory (0/39) and inhibitory (0/8) animal (per-animal incidence 3/5 control vs 0/9 DREADD, Fisher’s exact P = 0.027). Excitatory responses were almost entirely contralateral and selective (38/39), whereas inhibitory responses were as often bilateral as contralateral (4/8 bilateral). Local injection of muscimol (0.5 µL, 5 mM) into the ipsilesional cortex abolished evoked responses associated with the interhemispheric pathway in a representative excitatory animal. Sample sizes: n = 4–5 per group. Alt text: Intracortical microstimulation response maps of the left and right motor cortex for control, excitatory, and inhibitory animals, in which marker shape, size, and colour encode the evoked-movement type, threshold current, and side of movement; excitatory animals show an expanded, correctly lateralized right-forelimb representation.

We classified each contralesional forelimb response by the side of the evoked movement. In control animals, most responses drove the left (intact) rather than the right (impaired) forelimb (11 of 15 responses, in 3 of 5 animals), indicating maladaptive cortical re-lateralization away from the injured limb. This pure ipsilateral mistargeting was absent from every DREADD animal: it occurred in 3 of 5 control animals but none of the 9 excitatory or inhibitory animals (Fisher’s exact P = 0.027), and at the response level in 0 of 39 excitatory and 0 of 8 inhibitory responses (Fisher’s exact P < 0.0001 and P = 0.001 respectively, both vs. control). The groups differed in what they did instead: excitatory animals drove the right forelimb almost exclusively and selectively (38 of 39 responses purely contralateral, one bilateral), whereas the responses evoked in inhibitory animals were sparse and non-selective (four of eight were bilateral). Excitatory chemogenetic activity during play therefore preserved the correct lateralization and contralesional drive to the impaired forelimb.

At the animal level, the per-rat proportion of contralesional stimulations evoking a concordant (right) forelimb response differed significantly between groups (Kruskal–Wallis H = 9.08, exact P = 0.0031; Fig. 5). Dunn’s multiple comparisons confirmed a higher concordant proportion in excitatory animals than in both controls (adjusted P = 0.0236) and inhibitory animals (adjusted P = 0.0367), while control and inhibitory animals did not differ (P > 0.99). Median concordant proportions were 0.21 (excitatory), 0.02 (control) and 0.02 (inhibitory).

### The formation of corticospinal relay circuits bypassing the lesion

To determine how cortical activity influenced recruitment of descending relay pathways, we used an intersectional viral tracing strategy (Fig. 6A) in which BFP labels the retrogradely projecting spinal population and Cre-dependent mCherry is restricted to neurons that received transsynaptic input from left forelimb cortical neurons; the mCherry⁺ proportion thus indexes cortical relay neurons forming novel connections to the distal spinal cord. The extent of post-synaptic labelling did not differ between excitatory and control groups at any level. Reduction in the inhibitory group reached pairwise significance at the spinal cord (Ex vs. Inh p = 0.006; Con vs. Inh p = 0.033), consistent with AAV1 trans-synaptic transfer requiring neural activity.

**Figure 6.**
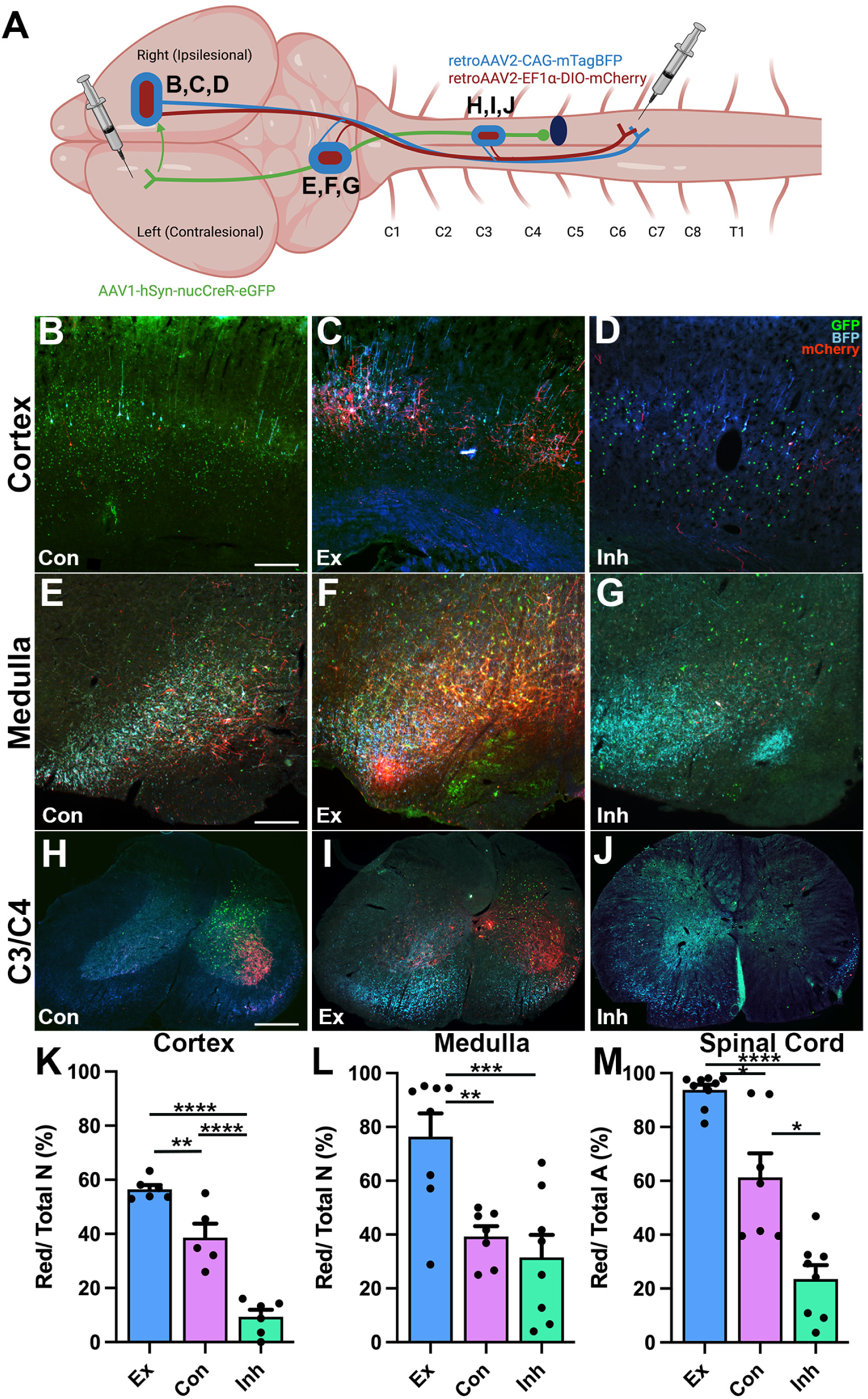
Intersectional viral tracing reveals chemogenetic excitation drives recruitment of interhemispheric, reticulo-spinal, and propriospinal bypass circuits. **(A)** Tracing strategy: AAV1-hSyn-nucCreR-eGFP injected into left (contralesional) motor cortex; 1:1 mixture of retrograde AAV2-mTagBFP (Cre-independent) and AAV2-DIO-mCherry (Cre-dependent) injected into right (ipsilesional) C6–T1 spinal cord. Green nuclei = AAV1 transsynaptic targets; blue = retrogradely labelled neurons; red = double-labelled cortical relay neurons. **(B–D)** Right (ipsilesional) primary motor cortex from control **(B)**, excitatory **(C)**, and inhibitory **(D)** animals. **(E–G)** Left (contralesional) Gi/LPGi medullary reticular nuclei from control **(E)**, excitatory **(F)**, and inhibitory **(G)** animals. **(H–J)** Right (ipsilesional) C3/C4 spinal cord from control **(H)**, excitatory **(I)**, and inhibitory **(J)** animals. Scale bars: 200 µm **(B–D)**; 500 µm **(E–G, H–J)**. Channel key shown in **(D)** applies to all microscopy panels. **(K–M)** Quantification of mCherry+ proportion at three anatomical levels; group means ± SD are reported in Results. **(K)** Ipsilesional cortex (n = 6 / 5 / 6 Ex / Con / Inh): %mCherry+ of total BFP+ neurons; one-way ANOVA F(2,14) = 56.96, p < 0.0001, ηZ = 0.89; Tukey’s: Ex–Con p = 0.0053, Ex–Inh p < 0.0001, Con–Inh p < 0.0001. **(L)** Contralesional medulla (n = 8 / 7 / 8): one-way ANOVA F(2,20) = 10.46, p = 0.0008, ηZ = 0.51; Tukey’s: Ex–Con p = 0.007, Ex–Inh p = 0.001, Con–Inh p = 0.76 (ns). **(M)** Ipsilesional C3/C4 (n = 9 / 7 / 8): %mCherry+ area relative to total mCherry+ + BFP+ area within spinal grey matter; Welch’s ANOVA W(2, 10.01) = 78.68, p < 0.0001, ηZ = 0.81; Games–Howell: Ex–Con p = 0.024, Ex–Inh p < 0.0001, Con–Inh p = 0.011. Bars represent mean ± SEM with individual data points overlaid; Excitatory = blue, Control = magenta, Inhibitory = teal. * p < 0.05, ** p < 0.01, *** p < 0.001, **** p < 0.0001. Alt text: An intersectional-tracing schematic with fluorescence micrographs of the cortex, medulla, and C3/C4 spinal cord for the three groups, plus bar graphs showing the highest proportion of double-labelled relay neurons in excitatory animals at all three levels.

In excitatory animals, we also observed a small population of ipsilesional (right) cortical neurons projecting to spinal cord caudal to the lesion. This was not attributable to incomplete lesioning, as histology confirmed complete hemisection at the epicentre (Fig. S5). The presence of Cre-dependent labelling in ipsilesional corticospinal populations is consistent with strengthened interhemispheric coupling and recruitment of intact ipsilesional CST neurons into compensatory circuits. Quantitative analysis revealed striking group differences in the percentage of ipsilesional (right) red-labelled cortical relay neurons across excitatory (56.5 ± 3.9%), control (38.8 ± 11.7%), and inhibitory (9.3 ± 6.4%) animals, with all pairwise contrasts significant (one-way ANOVA p < 0.0001; Fig. 6K). Overall total neuronal counts did not differ across groups (Fig. S6A), confirming that the relay-recruitment differences were not attributable to baseline neuronal number or survival.

To determine whether chemogenetic stimulation altered relay circuits at the brainstem level, we quantified labelled neurons in the contralesional gigantocellular (Gi) and lateral paragigantocellular (LPGi) nuclei (Fig. 6E–G) — reticulospinal populations that are important mediators of locomotor recovery after unilateral cervical hemisection.^5^ Quantification revealed a strong group effect in the percentage of red-labelled neurons in the contralesional (left) medulla (one-way ANOVA p = 0.0008; Fig. 6L), with excitatory animals exhibiting the highest proportion (76.4 ± 24.5%) compared with controls (39.3 ± 10.2%) and inhibitory animals (31.6 ± 23.5%). Overall total neuronal counts did not differ across groups (Fig. S6C).

To test whether intersectional relay recruitment extends to the spinal cord rostral to injury, we quantified mCherry+ and BFP+ labelled area in right-sided C3/C4 sections (ipsilesional to the P14 C5 hemisection; Fig. 6M). Consistent with the cortex and medulla, excitatory DREADD animals (93.8 ± 5.9%) exhibited the greatest red-labelled neuron coverage, whereas controls (61.3 ± 23.4%) were intermediate and inhibitory animals (23.6 ± 14.6%) showed sparse labelling. Welch’s ANOVA of red-labelled neuron coverage revealed a robust group effect (p < 0.0001). Post hoc comparisons confirmed that excitatory animals differed significantly from both control (p = 0.024) and inhibitory groups (p < 0.0001), and control exceeded inhibitory (p = 0.011). These results indicate that excitatory stimulation increased the overall recruitment of red-labelled neurons rostral to the lesion, consistent with reinforced cortico–propriospinal bypass circuits. Since the right dorsal and dorsolateral funiculus is destroyed at the C5 hemisection, these ipsilesional C3/C4 relay neurons likely access caudal spinal segments via axons descending through the spared ventral and ventromedial funiculus, consistent with the known trajectory of propriospinal projections through laminae VII and VIII.^1,2^

### Excitatory modulation rescues H-reflex spinal presynaptic inhibition

Maladaptive plasticity can lead to excessive spasticity or clonus due to decreased descending inhibition onto proprioceptive networks. Spasticity can also reduce rehabilitative success by restricting movements. To assess whether excitatory cortical stimulation influenced spinal inhibitory circuits, we examined frequency-dependent depression (FDD) of the H-reflex recorded from the ulnar nerve and abductor digiti minimi (ADM) muscle (n = 8 excitatory, n = 9 controls; Fig. 7A–B). In intact animals, repetitive stimulation produces progressive suppression of H-reflex amplitude at higher frequencies, reflecting normal presynaptic inhibition. After C5 hemisection, this suppression is diminished, indicating pathological hyperreflexia.

**Figure 7.**
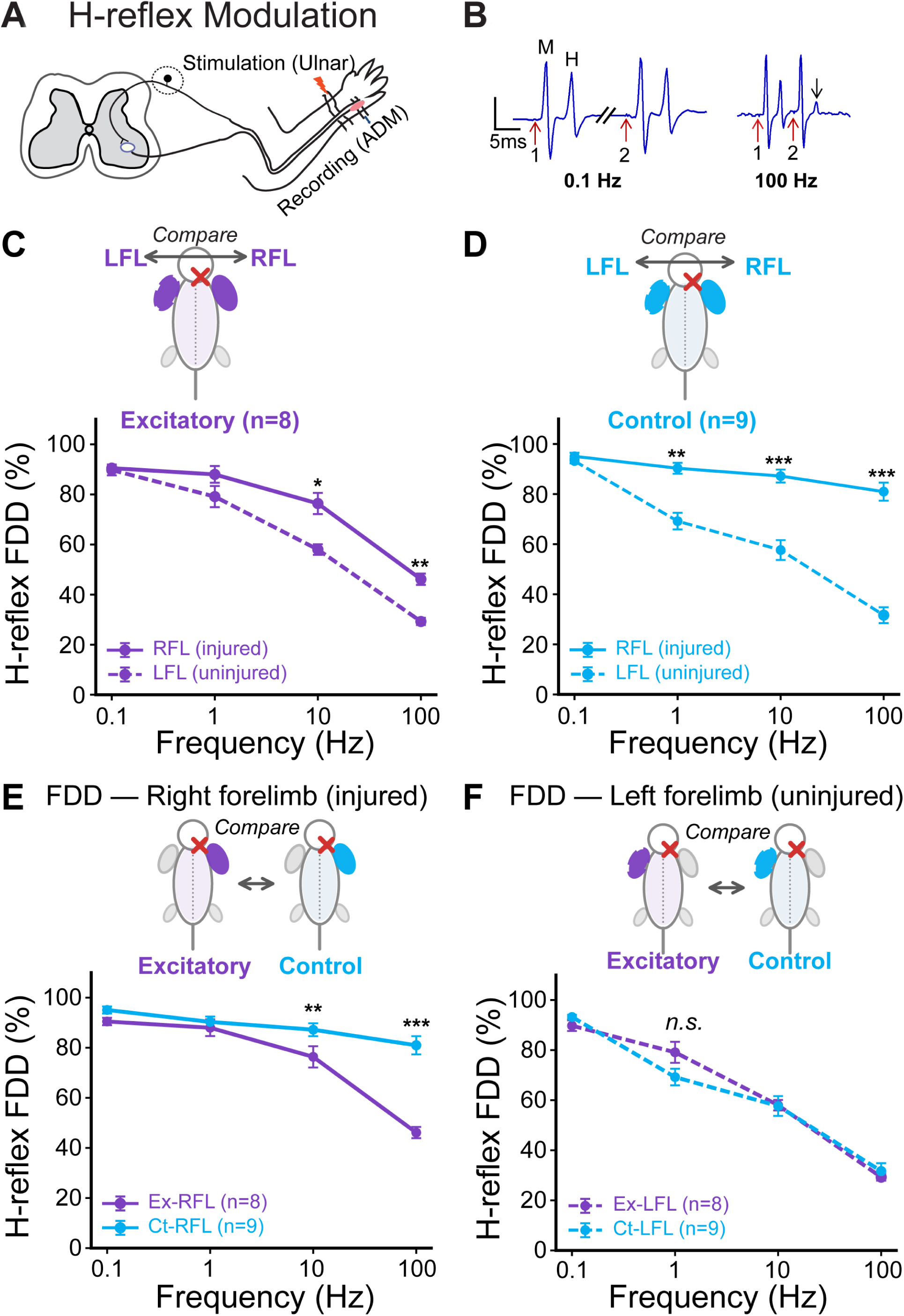
Excitatory chemogenetic stimulation restores frequency-dependent depression of the H-reflex in the injured forelimb. **(A)** Schematic of H-reflex testing. The ulnar nerve was stimulated at the wrist; EMG responses were recorded from the ipsilateral abductor digiti minimi (ADM). A conditioning–test paired-stimulus protocol assessed frequency-dependent depression (FDD) across 0.1–100 Hz. **(B)** Representative EMG traces showing intact FDD: at 100 Hz, the test H-reflex (arrow) is markedly suppressed relative to the conditioning H-reflex; at 0.1 Hz, both H-reflexes are similar in amplitude. **(C, D)** Within-group comparison of FDD between injured (right, RFL) and uninjured (left, LFL) forelimbs in excitatory **(C)** and control **(D)** animals. **(E, F)** Between-group comparison of injured **(E)** and uninjured **(F)** forelimbs across excitatory (n = 8) and control (n = 9) cohorts. Excitatory animals show restored high-frequency suppression of the injured-limb H-reflex (E; ∼46% at 100 Hz vs. ∼80% in controls), while uninjured-limb suppression is comparable across groups **(F)**, confirming lateralized therapeutic effect. Within-group comparisons **(C, D)**: two-way repeated-measures ANOVA (limb × frequency), with Bonferroni-corrected post-hoc. Excitatory: limb × frequency F(3,21) = 4.02, p = 0.042, ηZG = 0.18. Control: limb × frequency F(3,24) = 40.76, p < 0.001, ηZG = 0.53. Between-group comparisons **(E, F)**: two-way mixed ANOVA (group × frequency), Bonferroni-corrected post-hoc. Injured **(E)**: group F(1,15) = 35.60, p < 0.001, ηZp = 0.70; frequency F(3,45) = 44.74, p < 0.001, ηZp = 0.75; group × frequency F(3,45) = 15.65, p < 0.001, ηZp = 0.51. Uninjured **(F)**: group p = 0.65 (ns). Values are mean ± SEM. Alt text: A schematic of H-reflex testing with example EMG traces and frequency–response plots showing that excitatory animals restore high-frequency suppression (frequency-dependent depression) of the injured-forelimb H-reflex toward the level of the uninjured side, unlike control animals.

Consistent with this pathology, control animals showed significantly impaired FDD in the injured (right) forelimb compared with the intact (left) side (p < 0.001; Fig. 7D). At 100 Hz, the uninjured left forelimb retained robust suppression (31.7 ± 9.5% of baseline) whereas the injured right forelimb showed minimal depression (81.0 ± 10.9% of baseline), representing a ∼50 percentage-point difference (Cohen’s d = 3.1).

By contrast, excitatory DREADD-expressing rats exhibited marked restoration of FDD in the injured limb (Fig. 7E). A two-way mixed ANOVA comparing the injured (right) forelimb between excitatory and control groups confirmed significant main effects of frequency (p < 0.001) and group (p < 0.001), and a significant group × frequency interaction (p < 0.001; Fig. 7E). Post hoc comparisons revealed that excitatory animals showed significantly greater suppression than controls at 10 Hz (p = 0.009) with a ∼35 percentage-point greater suppression than controls at 100 Hz (p < 0.001). Effect sizes were large across all comparisons (ηZ > 0.4), underscoring the magnitude of recovery. Importantly, the lateralization of the therapeutic effect is reflected in the between-group effect-size contrast at 100 Hz: Cohen’s d = 3.86 for the injured (right) limb vs. d = 0.32 for the uninjured (left) limb, confirming that excitatory DREADDs improved FDD selectively in the injured limb.

Thus excitatory cortical activation reinstated frequency-dependent depression after paediatric SCI, indicating functional re-engagement of spinal inhibitory circuits (the inhibitory cohort was not recorded). This circuit-level normalization provides a mechanistic substrate for the superior functional recovery of the excitatory group.

### Convergent evidence for circuit reorganization and functional restoration

Across all experimental domains, a consistent hierarchy emerged: excitatory animals showed superior behavioural recovery, restored reaching kinematics, expanded contralesional cortical maps, increased recruitment of cortical, medullary, and spinal relay neurons into lesion-bypassing circuits, and reinstated frequency-dependent depression of the H-reflex. These convergent results demonstrate that enhancing cortical excitability during the juvenile plasticity window re-routes descending output through three parallel bypass circuits: (1) interhemispheric cortico–cortical–spinal connections, in which ipsilesional (right) cortical neurons project to the caudal spinal cord; (2) cortico–reticulo–spinal relays, in which contralesional (left) Gi and LPGi neurons receive ipsilateral corticobulbar drive and project across the spinal midline into the denervated hemicord; and (3) cortico–propriospinal relays, in which ipsilesional (right) C3/C4 interneurons receive crossed corticospinal input and project caudally past the lesion. Together, this reorganization partially restores motor representations and normalizes spinal reflex physiology.

## Discussion

The central aim of this study was to determine whether modulating cortical excitation during juvenile play altered descending motor systems and functional adaptability following SCI. Play is widely used in paediatric rehabilitation to boost engagement, yet is rarely studied mechanistically in SCI.^61–63^ In rodents, rough-and-tumble play provides patterned motor activity that stimulates sensorimotor circuits. We hypothesized that coupling chemogenetic excitation to behaviourally meaningful activity would direct plasticity towards adaptive circuit formation. Across all paradigms, goal-directed activity synergistically reinforced elevated excitability to consolidate adaptive circuitry.

A paradox in spinal cord research is that anatomical growth does not inexorably yield commensurate function: Gennaro et al.^11^ showed that P14 lesions produced greater axonal regrowth but worse adult motor outcomes than P21 lesions. Our findings address this gap directly: chemogenetic excitation paired with play steered nascent corticospinal and subcortical projections toward functional relays and drove significant behavioural gains, whereas reducing cortical activity dampened plasticity and yielded the poorest recovery.

Inducible excitation during the juvenile period likely reinforced activity-dependent consolidation across the motor pathway. Because patterned corticospinal activity promotes contralateral sprouting,^64,65^ repeatedly engaging corticospinal, reticulospinal, and propriospinal populations during play preserved cortical topography and favoured Hebbian stabilization of forelimb relays.^66,67^ Mirror-like activity within parieto-frontal circuits during social play may have further amplified this coordinated recruitment.^67^

The muscimol-induced abolition of contralesional responses supports interhemispheric co-activation through shared descending targets.^68–70^ Similar bilateral recruitment occurs in primates and humans after forelimb training, via dense callosal projections between motor cortices.^71^

In addition to interhemispheric recruitment, activity-dependent plasticity strengthened corticobulbar and propriospinal networks, selectively recruiting anatomically preserved neurons into lesion-bypassing relays.^65^ Engmann et al.^5^ established that spared contralesional Gi and LPGi neurons sprout into the denervated hemicord and that silencing them reimpairs locomotion; our data extend this to the juvenile cord, where modulating cortical excitation during a developmental plasticity window biased connectivity toward these same reticulospinal relays.

All injured animals advanced the shoulder (innervated above the C5 lesion) to compensate, but only excitatory animals achieved the wrist–shoulder separation to clear the opening and greater affected-forelimb use in the cylinder — a task requiring elbow extension against gravity. We interpret this as restoration of descending drive through reticulospinal relays onto C6–C8 motor pools governing elbow and wrist extension. The reticulospinal tract is known to preferentially target proximal and intermediate limb muscles, consistent with recovery of reach extension and cylinder exploration.^72–74^

H-reflex was assessed in the ulnar nerve and abductor digiti minimi, innervated by C8– T1 motor pools below the lesion. Restoration of FDD in excitatory animals therefore indicates that descending inhibitory tone was re-established distal to the injury. Partial restoration of fine forelimb function (improved IBB scores in excitatory animals) corroborated this. Restored inhibitory gating, coupled with incomplete digit manipulation, aligns with the reported limits of the cortico-reticulo-spinal pathway: the lateral reticulospinal tract restores inhibitory interneuron drive to distal cervical segments via laminae VII and VIII, but cannot fully substitute for the direct CST innervation required for independent finger control.^72,73^

The mechanisms we identified are consistent with those observed in spinal stimulation, where patterned epidural stimulation promotes locomotion by activating propriospinal afferents (V2a interneurons) and entrains spinal circuits during rehabilitation.^75^ Chemogenetic excitation during play functioned as a developmental analogue for epidural electrical stimulation (EES), integrating top-down cortical drive with meaningful activity. In contrast, continuous spinal stimulation absent of coordinated physical activity induces synaptic fatigue, asynchronous firing, and ultimately impairs functional recovery.^76^ In traumatic cervical SCI patients, the rate of functional motor improvement during inpatient rehabilitation is a stronger predictor of long-term independence than injury severity at admission.^77^ The therapeutic value of excitation lies in its timing, not magnitude; plasticity must be driven by behaviourally significant activity and not constitutive depolarization.

Conventional neuromodulation relies on implanted hardware, which is susceptible to impedance drift and hardware migration.^78^ Chemogenetic approaches offer pathway-specific, distributed modulation without implanted electrodes.^29^ Our results reinforce this: inducible excitation coupled with natural play produced multimodal descending reorganization, integrating regenerative biology with neuromodulation.

Animal models provide mechanistic resolution but cannot fully capture the heterogeneity and chronicity of human SCI. Our hemisection paradigm isolates descending disconnection and enables anatomical precision yet lacks the complex pathology of clinical contusions. In addition, the lesions were delivered early in development and thus were unable to test whether prehabilitation would impact recovery. Early-phase inhibition using pharmacologically selective actuator modules (PSAMs) in the first post-injury week, followed by excitatory upregulation, may delineate optimal temporal sequencing of neuronal modulation. Chemogenetic tools offer exquisite specificity but require clinically viable delivery strategies. A brain–spine interface that translates decoded cortical intention into spatiotemporal epidural stimulation has already restored naturalistic walking in chronic tetraplegia;^79^ integrating such closed-loop platforms with pathway-specific chemogenetic priming could enable multi-site control of recovery.

Recovery after paediatric cervical SCI hinges on which connections form, not simply how many. By aligning cortical excitation with natural play, we reconstituted diffuse sprouting into structured relay formation that restored forelimb movement. The broader message is clear: growth creates possibility and guided activity creates function.

## Data availability

Raw and processed kinematic data from the high-speed reaching task, together with the per-animal Source Data Excel workbook underlying the main and supplementary figures, are deposited at Figshare (https://doi.org/10.6084/m9.figshare.32335200). The trained DeepLabCut networks used to track wrist, elbow, and shoulder markers in high-speed video are deposited as a separate Figshare article (https://doi.org/10.6084/m9.figshare.32337243). All analysis code (Python scripts for kinematic feature extraction, SPM1d analyses, ICMS proportion and laterality categorization, and figure generation) is archived at Zenodo (https://doi.org/10.5281/zenodo.20309608).

## Acknowledgements

We would like to thank Drs. Sergio Pellis and Jackson Ham for discussions and methods on scoring play behaviour. We are also grateful to the many undergraduate students for their dedicated assistance over the years.

## Supplementary material

Supplementary material accompanies this preprint. This includes Supplementary Methods, Supplementary Figures S1–S8, Supplementary Tables S1–S5, and Supplementary Video 1.

