## Supplementary Methods, Figures S1-S8, Tables S1-S5 for "Chemogenetic excitation and play promote functional recovery in juvenile spinal cord injury"

*Viral vector production.* HEK293T cells (ATCC CRL-3216) at 70–80% confluency were transfected (polyethylenimine, linear MW 25 kDa, 1:3 DNA:PEI) with the transgene, an AAV rep/cap packaging plasmid, and an adenoviral helper plasmid. After 72 h, crude virus was precipitated with 40% PEG 8000 (4 °C, 2 h), resuspended in 10 mM Tris–HCl (pH 8.0), purified by double caesium-chloride ultracentrifugation, dialysed against 5% sorbitol/PBS, and stored at −80 °C.

*Kinematic acquisition and analysis.* Reaches were made through a 10 mm vertical slit in a transparent acrylic chamber (100 × 50 × 50 mm), pellet 30 mm beyond the wall, recorded at 250 frames $s^{-1}$ with a Ximea XiQ USB3 high-speed camera (XIMEA GmbH) under infrared backlighting, camera and lens fixed across animals. Shoulder, elbow, wrist and digit-tip markers were labelled in ~200 frames per view and tracked with DeepLabCut v2.3 (ResNet-50 backbone, 500,000 iterations, standard augmentation, p-cutoff 0.9). For SPM analysis, shoulder, elbow, and wrist time series were aligned to the reach-event peak frame and resampled to 200 points across the peri-reach interval; tracking dropouts were linearly interpolated within each reach, trajectories truncated to a common window, and coordinates converted to millimetres using a session-specific calibration. Inhibitory elbow trajectories were low-pass filtered (3rd-order Butterworth, 4–6 Hz) per reach; no other condition was filtered.

*ICMS stimulation parameters.* A tungsten microelectrode (0.5 MΩ; FHC) was advanced perpendicular to the cortex to 1.8 mm (layer V) with a hydraulic micropositioner; biphasic pulses (0.2 ms per phase, cathodal-leading) were delivered in 300 ms trains at 100 Hz.

*Intersectional tracing coordinates.* Cortical AAV1-hSyn-nucCreR-eGFP: 100 nL $min^{-1}$ across four sites (2 µL per site, 8 µL total; from bregma AP +2.0, ML –1.4; AP +2.0, ML –2.6; AP +1.0, ML –1.4; AP +1.0, ML –2.2 mm; depth 1.5 mm), pipette held 5 min before withdrawal. Spinal retrograde mixture: ten sites along C6–T1 (300 nL per site, 3 µL total; ML +0.8 mm, depth 1.5 mm from dorsal surface), intermediate grey matter.

*H-reflex recording.* The ulnar nerve was stimulated at the wrist via paired subcutaneous braided-wire electrodes (0.014 mm; AS 634, Cooner Wire) using a constant-current isolated stimulator (100 µs pulse; A–M Systems); EMG was recorded from abductor digiti minimi via two intramuscular fine-wire electrodes.

*Neonatal cortical injections.* Viral suspension (0.5 µL per site at 100 nL $min^{-1}$) was delivered via bevelled glass micropipettes (tip 20–30 µm) to the left forelimb motor cortex at three sites (coordinates from bregma: AP +0.5 mm; ML –1.0, –1.5, and –2.0 mm; depth –0.6 mm from the cortical surface), with the pipette retained in situ for 3 min before withdrawal.

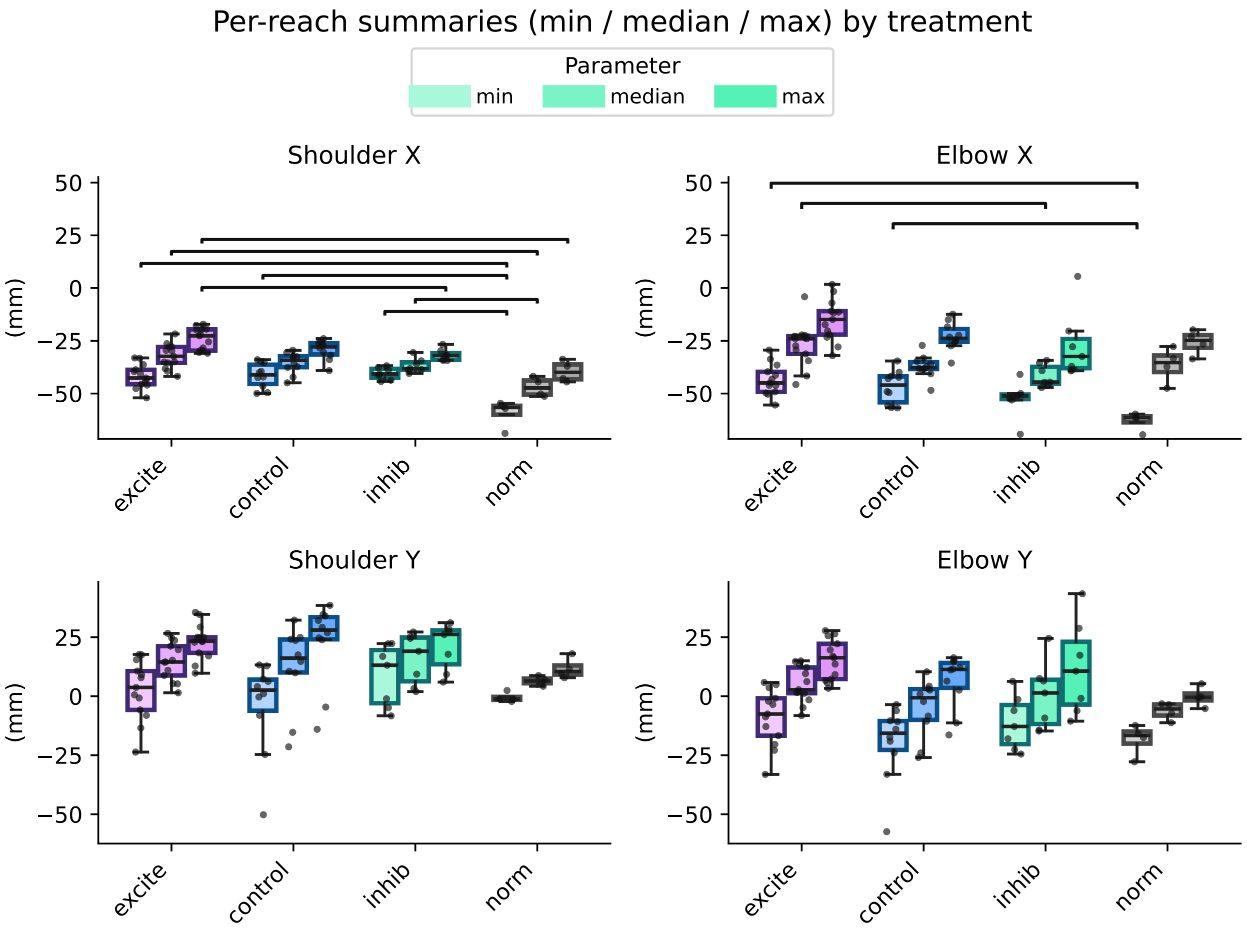


**Figure S1.** Variation in shoulder and elbow extremal kinematics with treatment. Excitatory animals bring their shoulder and elbow joints closer horizontally to the wall and pellet than inhibitory and normal (naïve, uninjured) animals, but the vertical position of these joints is not significantly different across groups. This may be a compensatory strategy that excitatory animals are able to perform that control and inhibitory injured animals cannot. Minimum, median, and maximum values of shoulder and elbow X (horizontal; toward the wall and pellet, +X; away from the wall, −X) and Y (vertical; upward, +Y; downward, −Y) coordinates are shown across treatment groups. Colours denote treatment (purple: excitatory; blue: control; green: inhibitory; grey: normal). Within each treatment, saturation indicates the summary parameter (minimum, median, maximum; see legend). Control animals were injured but untreated; normal animals were naïve and uninjured. Each point represents a single reach; boxplots summarize distributions across reaches. Group differences were assessed independently for each coordinate and summary parameter using Kruskal–Wallis nonparametric tests. Where significant omnibus effects were detected ($p < 0.05$), pairwise Mann–Whitney U tests with Holm correction were performed. Horizontal overbars indicate statistically significant pairwise comparisons. Omnibus statistics in Table S2; full pairwise comparisons in Table S3, with significant comparisons (after Holm correction) shown as overbars in this figure. n_reaches = 13 / 10 / 7 / 4 (Excitatory / Control / Inhibitory / Normal).

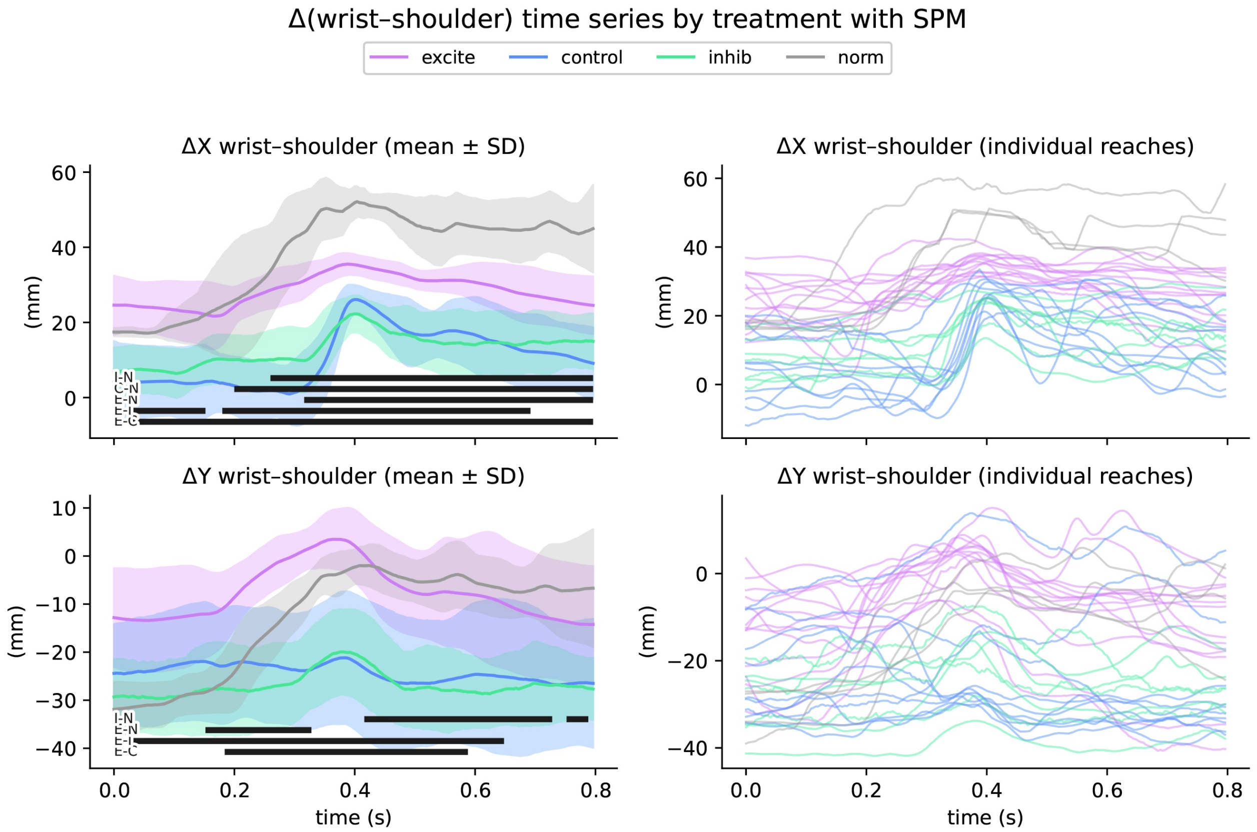


**Figure S2.** Wrist–shoulder relative kinematics across treatments. Time-varying wrist position relative to the shoulder is shown as ΔX (horizontal; top row) and ΔY (vertical; bottom row), computed as wrist minus shoulder coordinates for each reach and expressed in mm. Left panels show mean ± SD trajectories for each treatment group; right panels show individual reaches. Colours denote treatment (purple: excitatory; blue: control; green: inhibitory; grey: normal). Pairwise differences between treatments were assessed using statistical parametric mapping (SPM; two-sample t-tests across the time-normalized trajectory), and significant clusters ($p < 0.05$, two-tailed) are indicated by horizontal bars near the x-axis. Pairwise comparisons indicated by letter codes: I–N inhibitory vs. normal; C–N control vs. normal; E–N excitatory vs. normal; E–I excitatory vs. inhibitory; E–C excitatory vs. control. Excitatory animals maintained a larger ΔX and ΔY separation between wrist and shoulder throughout the reach trajectory compared with control and inhibitory animals, indicating that excitatory animals successfully extended and elevated the wrist relative to the shoulder to clear the opening. Full SPM results (cluster time ranges and p-values) are provided in Table S1.

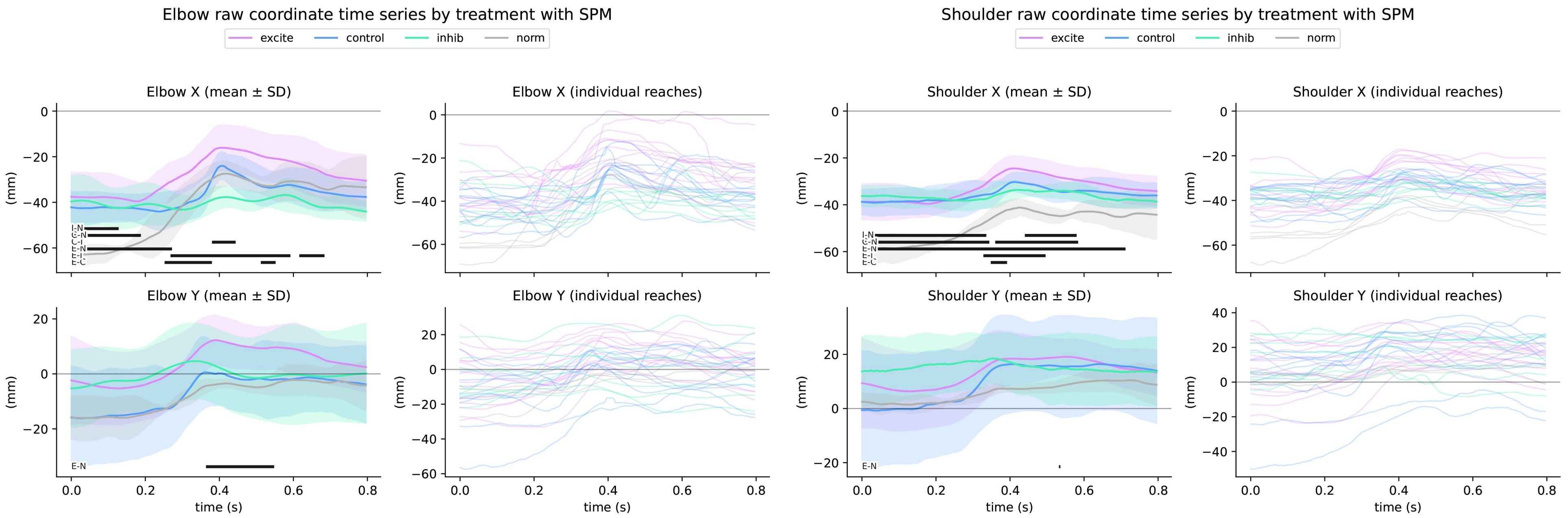


**Figure S3.** Elbow and shoulder raw coordinate kinematics across treatments. Time-varying elbow (left two columns) and shoulder (right two columns) X and Y coordinates are shown across treatment groups. X coordinates reflect horizontal position relative to the pellet and wall coordinate system (+X toward pellet, −X away); Y coordinates reflect vertical position (+Y upward, −Y downward). For each joint, the first column shows mean ± SD trajectories and the second column shows individual reaches. Colours denote treatment (purple: excitatory; blue: control; green: inhibitory; grey: normal). Inhibitory elbow trajectories were low-pass filtered to reduce tracking discontinuities before plotting and SPM analysis. Pairwise treatment differences were assessed using SPM two-sample t-tests across the trajectory, and significant clusters ($p < 0.05$, two-tailed) are indicated by horizontal bars near the x-axis. Pairwise comparisons indicated by letter codes: I–N inhibitory vs. normal; C–N control vs. normal; E–N excitatory vs. normal; E–I excitatory vs. inhibitory; E–C excitatory vs. control. All injured groups start closer to the pellet in elbow X (upper left), whereas uninjured normals start further from the pellet but retain the range of motion to reach fully to it; within injured groups, excitatory animals reach furthest in X, controls in the middle, and inhibitory animals are essentially unable to progress. For elbow Y, only the height difference between excitatory and normal animals reaches significance at peak reach. The shoulder data follow a similar pattern: in X, normal animals start further from the pellet yet reach to it; injured animals of all groups start closer to the pellet, and within those, excitatory animals reach furthest, then controls, then inhibitory. In shoulder Y, the data are noisier and no large regions of significance emerge, although similar trends to X are seen. Full SPM results in Table S4 (elbow) and Table S5 (shoulder).

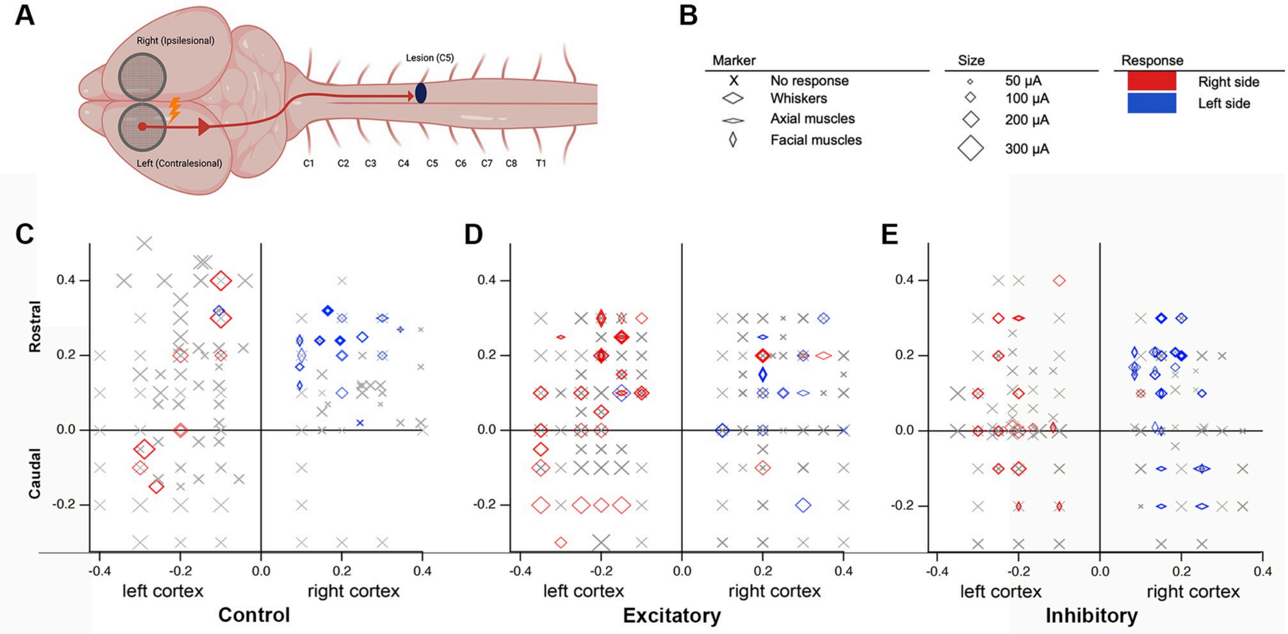


**Figure S4.** Intracortical microstimulation maps for non-forelimb body regions. Companion to Fig. 5, showing whisker, axial-muscle, and facial responses across the same stimulation grid in the same animals. (A) Schematic of bilateral cortical stimulation grid relative to the C5 lesion. (B) Marker key indicating responsive body region (whiskers = wide horizontal diamond, axial muscles = narrow horizontal diamond, facial muscles = narrow vertical diamond, no response = cross), threshold current encoded by marker size (50, 100, 200, or 300 µA), and side of evoked movement (red = right, blue = left). (C-E) Representative non-forelimb response maps for control (C), excitatory (D), and inhibitory (E) animals across left (contralesional) and right (ipsilesional) cortex; rostro-caudal grid position (mm relative to bregma) on the y-axis, mediolateral position on the x-axis. Excitatory animals (D) show widespread re-emergence of low-threshold whisker and axial sites in the contralesional cortex, paralleling the forelimb-domain effect shown in Fig. 5E and confirming that chemogenetic modulation alters cortical excitability across body-map representations rather than selectively in the forelimb domain. Sample sizes: n = 4–5 rats per group (same animals as Fig. 5).

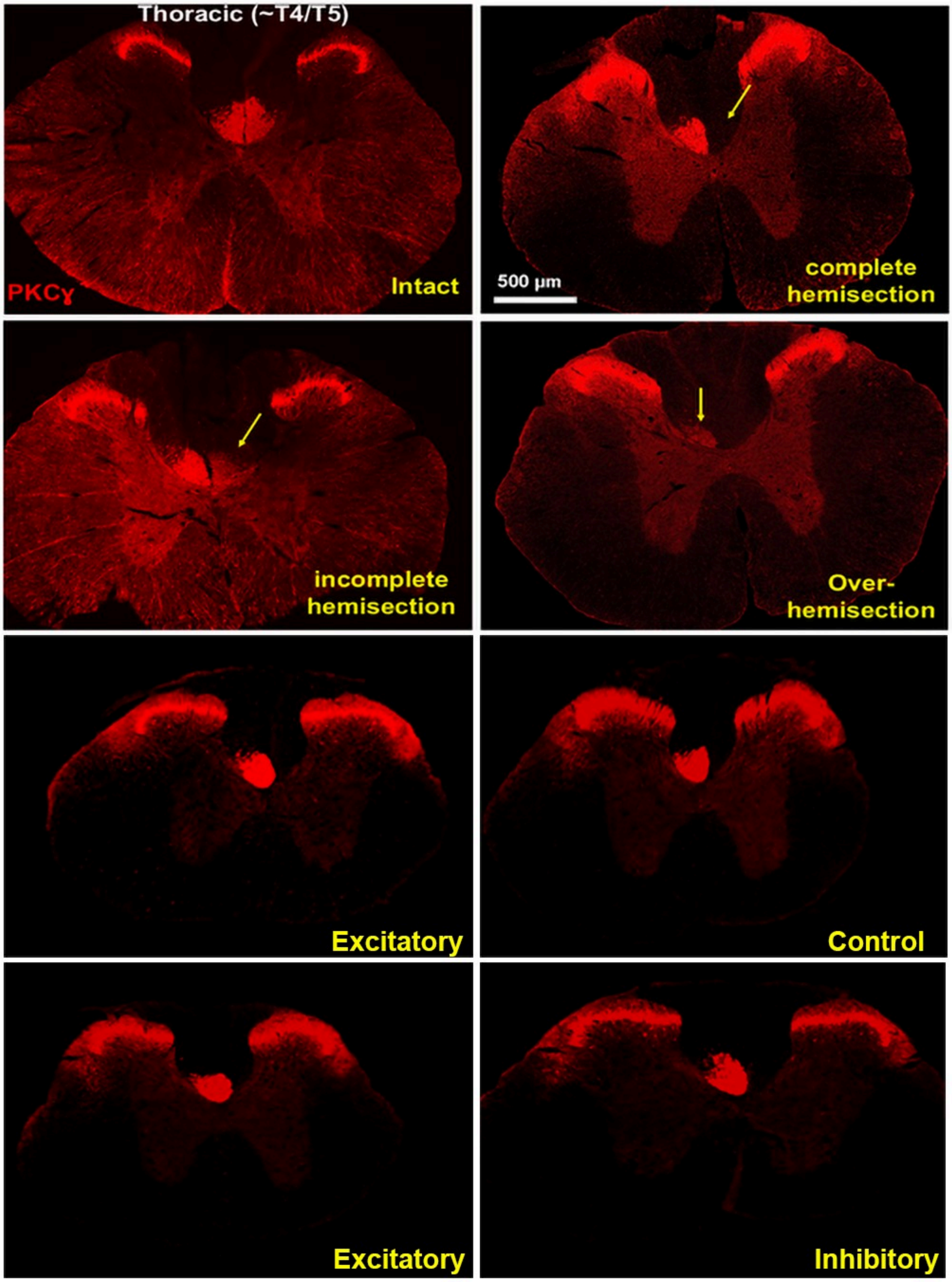


**Figure S5.** PKCγ verification of C5 hemisection completeness. Representative thoracic (~T4/T5) cross-sections immunolabelled for PKCγ (red), used to verify the integrity of corticospinal tract interruption rostral to the lesion. Top row, left: intact cord showing bilateral PKCγ-positive corticospinal tract in the dorsal funiculus. Top row, right: complete C5 hemisection with absence of PKCγ-positive fibres in the right (ipsilesional) dorsal funiculus (yellow arrow). Second row, left: incomplete hemisection showing residual ipsilesional PKCγ-positive fibres (yellow arrow); animals with this profile were excluded from analysis. Second row, right: over-hemisection with bilateral PKCγ loss (yellow arrow); also excluded. Third and fourth rows: representative complete-hemisection animals from the excitatory, mCherry control, and inhibitory cohorts retained for analysis, all showing intact contralesional and absent ipsilesional PKCγ labelling. Scale bar 500 µm. PKCγ: rabbit monoclonal, Abcam ab317315 (clone EPR28643-68), 1:200.

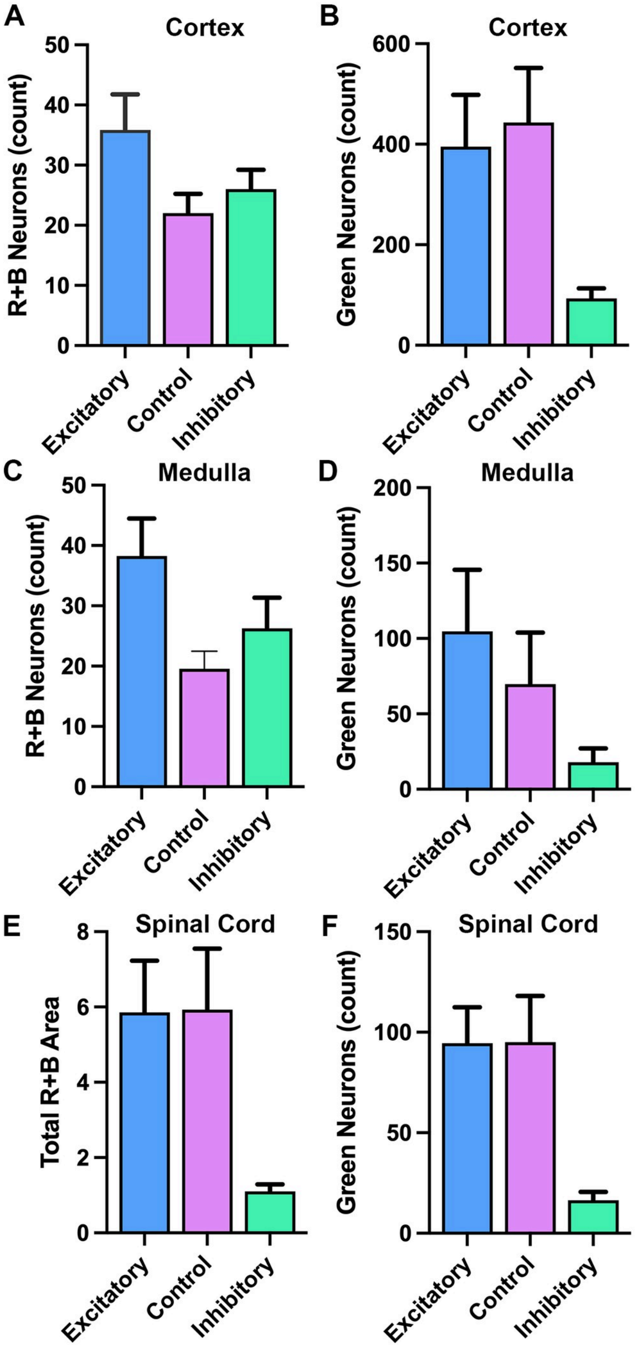


**Figure S6.** Total retrogradely-labelled (R+B) and post-synaptic (Green) neuron counts across cortex, medulla, and spinal cord. Companion to Fig. 6, providing absolute neuron counts (rather than the ratios shown in Fig. 6K–M) to assess whether the group differences in mCherry+/BFP+ percentages reported in the main figure are driven by overall differences in retrograde labelling efficiency or AAV1 trans-synaptic labelling. (A, C, E) Total counts of retrogradely BFP+ and mCherry+ neurons (R+B) in ipsilesional cortex (A; n = 6 / 5 / 6 Excitatory / Control / Inhibitory; one-way ANOVA $F(2,14) = 2.55$, $p = 0.113$, ns), contralesional medulla (C; n = 8 / 7 / 8; one-way ANOVA $F(2,20) = 3.40$, $p = 0.054$, ns), and ipsilesional C3/C4 spinal cord (E; n = 9 / 7 / 8; total mCherry+ + BFP+ labelled area in arbitrary units owing to dense, non-discrete propriospinal labelling; Welch's ANOVA $W(2, 9.44) = 9.45$, $p = 0.006$, with the inhibitory group reduced relative to excitatory and control, which were comparable). Cortex and medulla R+B counts therefore do not differ across groups; the reduction in spinal R+B area in the inhibitory group is acknowledged and is consistent with reduced overall propriospinal labelling rather than altered relay-neuron proportions, since the Fig. 6M ratio is normalized by total labelled area. (B, D, F) Total counts of post-synaptic AAV1-Cre-driven Green neurons in the same regions: ipsilesional cortex (B; n = 6 / 5 / 6; Welch's ANOVA $W(2, 6.33) = 8.01$, $p = 0.018$), contralesional medulla (D; n = 8 / 7 / 8; Welch's ANOVA $W(2, 9.59) = 2.87$, $p = 0.106$, ns), and ipsilesional C3/C4 spinal cord (F; n = 9 / 7 / 8; Welch's ANOVA $W(2, 9.68) = 13.57$, $p = 0.002$, with Games–Howell pairwise: Excitatory vs. Inhibitory $p = 0.006$; Control vs. Inhibitory $p = 0.033$; Excitatory vs. Control $p > 0.99$). Bars show mean ± SEM; individual data points overlaid. Excitatory = blue, mCherry control = magenta, Inhibitory = teal. The reduction in Green counts in the inhibitory group at the cortical and spinal levels (and a comparable trend at the medullary level) is consistent with AAV1 trans-synaptic transfer requiring intact neuronal activity.

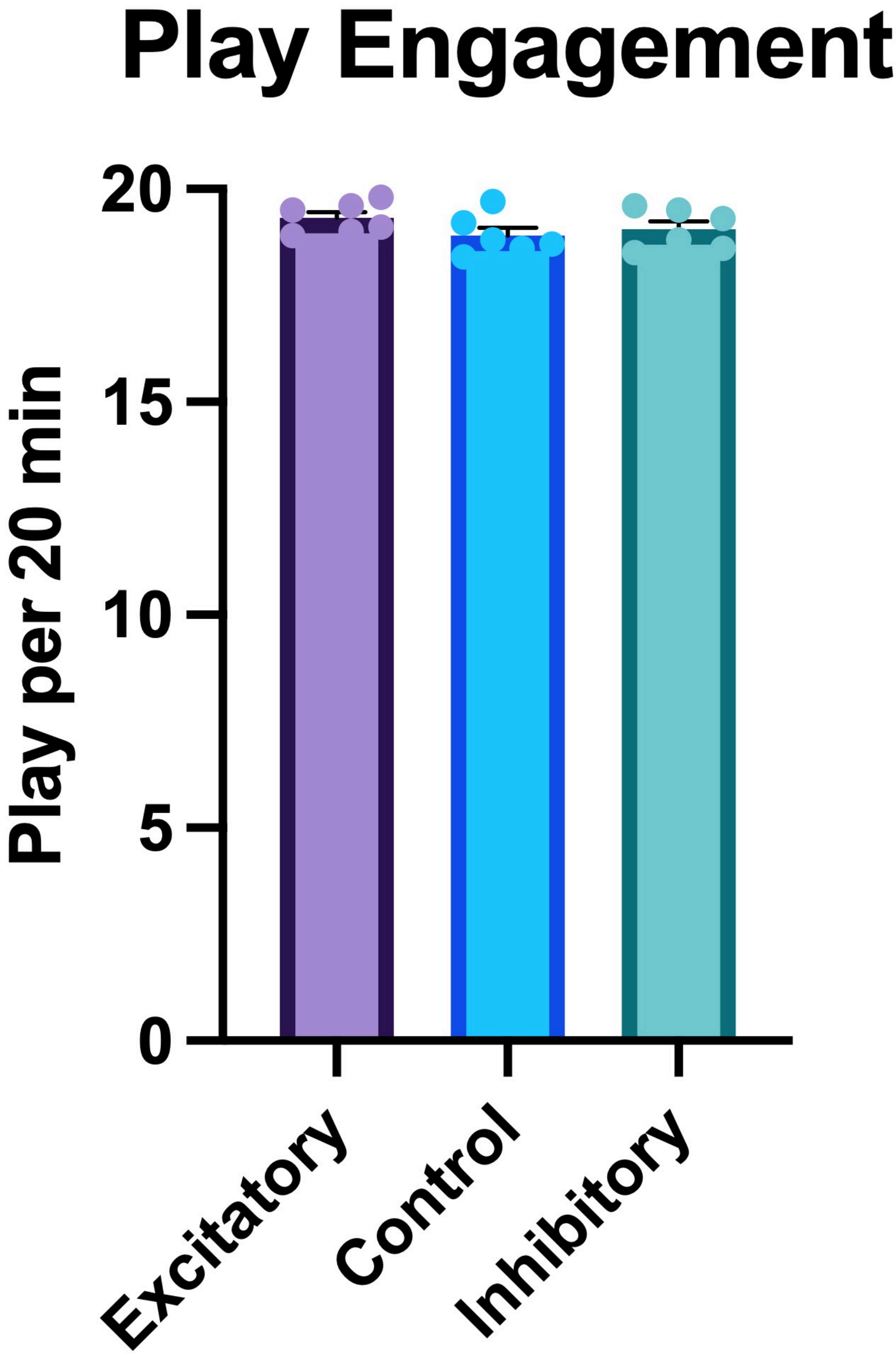


**Figure S7.** Chemogenetic group assignment does not alter play engagement among injured cohorts. To exclude the possibility that group differences in functional recovery (Fig. 3) reflected differential rough-and-tumble play participation under DREADD actuation, total time spent engaged in play was quantified during 20-minute nightly observation sessions across the P20–P50 CNO actuation window. All animals had received cortical AAV at P5 (Excitatory: AAV2-hSyn-hM3Dq-mCherry; Control: AAV2-hSyn-mCherry; Inhibitory: AAV2-hSyn-hM4Di-mCherry), undergone right C5 hemisection at P14, and were group-housed in cages of three same-sex littermates with nightly oral CNO from P20–P50, identical to the conditions of all behavioural and anatomical cohorts in this study. Bars show mean ± SEM with individual animal data points overlaid; Excitatory = purple ($n = 6$), Control = blue ($n = 6$), Inhibitory = teal ($n = 6$). One-way ANOVA confirmed no group difference in play engagement ($F(2,15) = 1.371$, $p = 0.2839$, $\eta^2 = 0.155$; means: Excitatory 19.32, Control 18.90, Inhibitory 19.05 minutes per 20-minute session) with no evidence of variance heterogeneity (Brown–Forsythe $p = 0.7141$; Bartlett $p = 0.8221$).

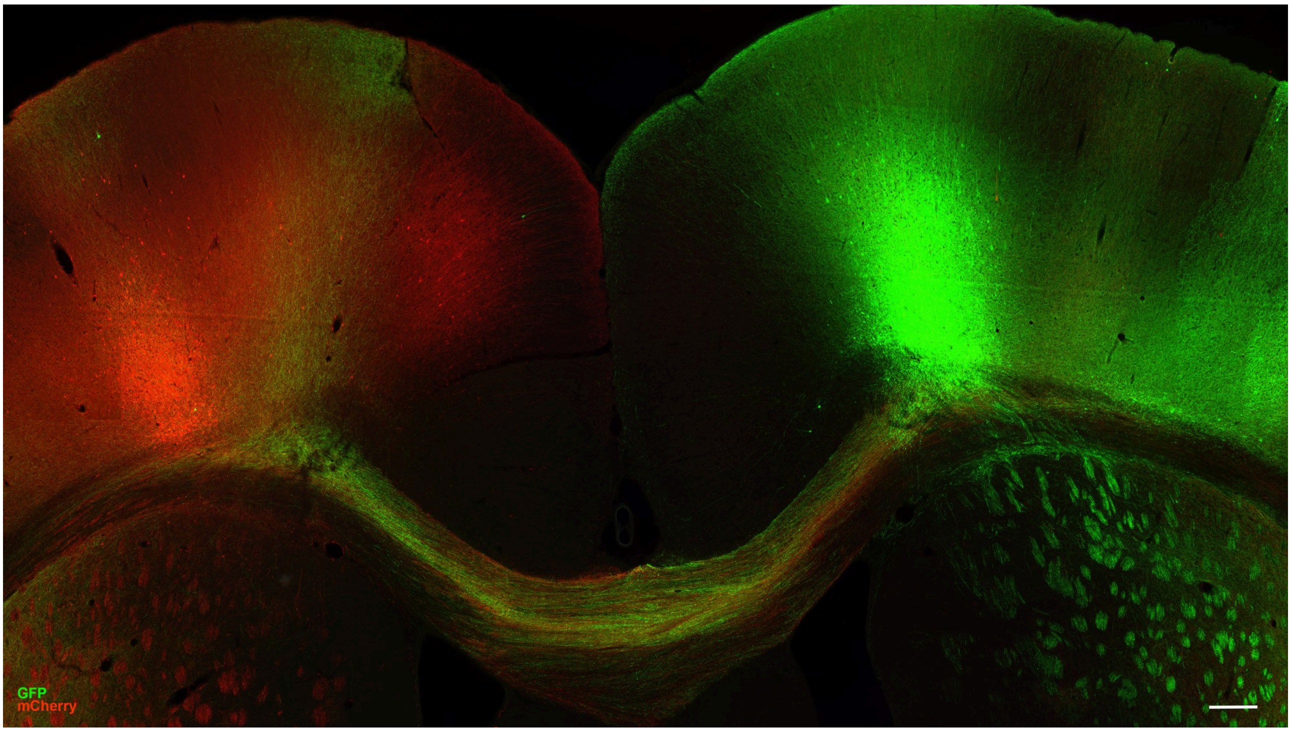


**Figure S8.** Bilateral cortical tracing reveals dense callosal projections between forelimb motor cortices in an excitatory animal. Representative coronal section through bilateral primary motor cortex from a single excitatory DREADD-treated animal that received AAV2-mCherry into the left (contralesional) forelimb motor cortex and AAV2-eGFP into the right (ipsilesional) forelimb motor cortex four weeks prior to perfusion. mCherry-labelled neurons (red) are visible at the left cortical injection site, GFP-labelled neurons (green) at the right cortical injection site, and labelled fibres from both populations cross and intermingle within the corpus callosum (yellow-orange region), demonstrating bidirectional homotopic callosal projections between the two M1 hemispheres. This anatomical connectivity provides a structural substrate consistent with the interhemispheric functional recruitment observed by ICMS in excitatory animals (Fig. 5; Fisher's exact P = 0.005 for ipsilesional cortex recruitment of right-forelimb evoked sites; abolition of evoked responses by ipsilesional muscimol injection). Tracing was performed as a qualitative anatomical complement to the quantified ICMS findings; n = 1 representative animal, no quantification was performed. Scale bar 1 mm.

**Table S1.** Significant SPM clusters for wrist–shoulder relative kinematics. Pairwise SPM t-tests identifying time intervals with significant differences in ΔX and ΔY wrist–shoulder trajectories between treatment groups ($p < 0.05$, two-tailed). Reported values include cluster time ranges, durations, and number of reaches per group. Companion to Fig. S2.

| measure | comparison | n1 | n2 | p | t_start | t_end | duration |
|---|---|---|---|---|---|---|---|
| ΔX wrist–shoulder | excite vs control | 13 | 10 | 0 | 0 | 0.796 | 0.796 |
| ΔX wrist–shoulder | excite vs inhib | 13 | 7 | 0 | 0.18 | 0.692 | 0.512 |
| ΔX wrist–shoulder | excite vs norm | 13 | 4 | 0 | 0.316 | 0.796 | 0.48 |
| ΔX wrist–shoulder | control vs norm | 10 | 4 | 0 | 0.2 | 0.796 | 0.596 |
| ΔX wrist–shoulder | inhib vs norm | 7 | 4 | 0 | 0.26 | 0.796 | 0.536 |
| ΔX wrist–shoulder | excite vs inhib | 13 | 7 | 7.333e-05 | 0 | 0.152 | 0.152 |
| ΔY wrist–shoulder | excite vs inhib | 13 | 7 | 0 | 0 | 0.648 | 0.648 |
| ΔY wrist–shoulder | excite vs norm | 13 | 4 | 0.0002054 | 0.152 | 0.328 | 0.176 |
| ΔY wrist–shoulder | inhib vs norm | 7 | 4 | 0.04004 | 0.752 | 0.788 | 0.036 |
| ΔY wrist–shoulder | excite vs norm | 13 | 4 | 0.0494 | 0 | 0.008 | 0.008 |
| ΔY wrist–shoulder | inhib vs norm | 7 | 4 | 4.417e-10 | 0.416 | 0.728 | 0.312 |
| ΔY wrist–shoulder | excite vs control | 13 | 10 | 6.749e-07 | 0.184 | 0.588 | 0.404 |

n1/n2 denote number of reaches in each group used for the SPM test; duration = t_end – t_start.

**Table S2.** Nonparametric omnibus tests for extremal shoulder and elbow kinematics. Summary of Kruskal–Wallis tests comparing treatment groups for minimum, median, and maximum shoulder and elbow kinematic coordinates (X and Y). Tests were conducted independently for each coordinate and summary parameter. Reported values include the Kruskal–Wallis H statistic and uncorrected p-values. The number of reaches contributing to each comparison is reported per treatment in the N (E/C/I/N) column. Companion to Fig. S1.

| coord | stat | N (E/C/I/N) | H | p_kw |
|---|---|---|---|---|
| shoulder_x | min | 13 / 10 / 7 / 4 | 10.548 | 0.01444 |
| shoulder_x | median | 13 / 10 / 7 / 4 | 12.248 | 0.00658 |
| shoulder_x | max | 13 / 10 / 7 / 4 | 17.431 | 0.0005761 |
| elbow_x | min | 13 / 10 / 7 / 4 | 13.802 | 0.003187 |
| elbow_x | median | 13 / 10 / 7 / 4 | 12.521 | 0.005797 |
| elbow_x | max | 13 / 10 / 7 / 4 | 8.410 | 0.03826 |
| shoulder_y | min | 13 / 10 / 7 / 4 | 2.371 | 0.499 |
| shoulder_y | median | 13 / 10 / 7 / 4 | 3.407 | 0.333 |
| shoulder_y | max | 13 / 10 / 7 / 4 | 5.426 | 0.1431 |
| elbow_y | min | 13 / 10 / 7 / 4 | 4.075 | 0.2535 |
| elbow_y | median | 13 / 10 / 7 / 4 | 6.483 | 0.09035 |
| elbow_y | max | 13 / 10 / 7 / 4 | 7.352 | 0.06147 |

H: Kruskal–Wallis test statistic. p_kw: uncorrected p-value.

**Table S3.** All pairwise post hoc comparisons for extremal kinematic measures. Results of pairwise Mann–Whitney U tests for every treatment comparison following each significant Kruskal–Wallis omnibus test in Table S2. For each comparison, the parent Kruskal–Wallis p-value, raw and Holm-corrected pairwise p-values, and significance level are reported. Sample sizes reflect the number of reaches per treatment contributing to each test. Provided as the full record of the pairwise structure underlying Fig. S1.

| coord | stat | comparison | N_A | N_B | KW_p | p_raw | p_adj | sig |
|---|---|---|---|---|---|---|---|---|
| shoulder_x | min | excite vs control | 13 | 10 | 0.01444 | 0.8768 | 1 | ns |
| shoulder_x | min | excite vs inhib | 13 | 7 | 0.01444 | 0.5356 | 1 | ns |
| shoulder_x | min | excite vs norm | 13 | 4 | 0.01444 | 0.0008403 | 0.005042 | ** |
| shoulder_x | min | control vs inhib | 10 | 7 | 0.01444 | 0.8868 | 1 | ns |
| shoulder_x | min | control vs norm | 10 | 4 | 0.01444 | 0.001998 | 0.00999 | ** |
| shoulder_x | min | inhib vs norm | 7 | 4 | 0.01444 | 0.006061 | 0.02424 | * |
| shoulder_x | median | excite vs control | 13 | 10 | 0.00658 | 0.2036 | 0.4072 | ns |
| shoulder_x | median | excite vs inhib | 13 | 7 | 0.00658 | 0.0968 | 0.2904 | ns |
| shoulder_x | median | excite vs norm | 13 | 4 | 0.00658 | 0.0008403 | 0.005042 | ** |
| shoulder_x | median | control vs inhib | 10 | 7 | 0.00658 | 0.4173 | 0.4173 | ns |
| shoulder_x | median | control vs norm | 10 | 4 | 0.00658 | 0.01399 | 0.05594 | ns |
| shoulder_x | median | inhib vs norm | 7 | 4 | 0.00658 | 0.006061 | 0.0303 | * |
| shoulder_x | max | excite vs control | 13 | 10 | 0.0005761 | 0.05075 | 0.1273 | ns |
| shoulder_x | max | excite vs inhib | 13 | 7 | 0.0005761 | 0.001161 | 0.005805 | ** |
| shoulder_x | max | excite vs norm | 13 | 4 | 0.0005761 | 0.0008403 | 0.005042 | ** |
| shoulder_x | max | control vs inhib | 10 | 7 | 0.0005761 | 0.1088 | 0.1273 | ns |
| shoulder_x | max | control vs norm | 10 | 4 | 0.0005761 | 0.01399 | 0.05594 | ns |
| shoulder_x | max | inhib vs norm | 7 | 4 | 0.0005761 | 0.04242 | 0.1273 | ns |
| elbow_x | min | excite vs control | 13 | 10 | 0.003187 | 0.4025 | 0.5397 | ns |
| elbow_x | min | excite vs inhib | 13 | 7 | 0.003187 | 0.01447 | 0.05789 | ns |
| elbow_x | min | excite vs norm | 13 | 4 | 0.003187 | 0.0008403 | 0.005042 | ** |
| elbow_x | min | control vs inhib | 10 | 7 | 0.003187 | 0.2698 | 0.5397 | ns |
| elbow_x | min | control vs norm | 10 | 4 | 0.003187 | 0.001998 | 0.00999 | ** |
| elbow_x | min | inhib vs norm | 7 | 4 | 0.003187 | 0.04242 | 0.1273 | ns |
| elbow_x | median | excite vs control | 13 | 10 | 0.005797 | 0.01006 | 0.05031 | ns |

| coord | stat | comparison | N_A | N_B | KW_p | p_raw | p_adj | sig |
|---|---|---|---|---|---|---|---|---|
| elbow_x | median | excite vs inhib | 13 | 7 | 0.005797 | 0.002425 | 0.01455 | * |
| elbow_x | median | excite vs norm | 13 | 4 | 0.005797 | 0.07899 | 0.316 | ns |
| elbow_x | median | control vs inhib | 10 | 7 | 0.005797 | 0.1932 | 0.5797 | ns |
| elbow_x | median | control vs norm | 10 | 4 | 0.005797 | 0.6354 | 0.8242 | ns |
| elbow_x | median | inhib vs norm | 7 | 4 | 0.005797 | 0.4121 | 0.8242 | ns |
| elbow_x | max | excite vs control | 13 | 10 | 0.03826 | 0.04385 | 0.2217 | ns |
| elbow_x | max | excite vs inhib | 13 | 7 | 0.03826 | 0.03695 | 0.2217 | ns |
| elbow_x | max | excite vs norm | 13 | 4 | 0.03826 | 0.07899 | 0.316 | ns |
| elbow_x | max | control vs inhib | 10 | 7 | 0.03826 | 0.1331 | 0.3992 | ns |
| elbow_x | max | control vs norm | 10 | 4 | 0.03826 | 0.6354 | 0.8242 | ns |
| elbow_x | max | inhib vs norm | 7 | 4 | 0.03826 | 0.4121 | 0.8242 | ns |

N_A and N_B denote the number of reaches in each comparison group. p_raw indicates the uncorrected pairwise Mann–Whitney U p-value; p_adj indicates the Holm-corrected p-value. KW_p indicates the corresponding Kruskal–Wallis omnibus p-value for that coordinate and summary statistic.

**Table S4.** Significant SPM clusters for elbow raw coordinate trajectories. Pairwise statistical parametric mapping analyses comparing treatment groups for time-varying elbow X and Y coordinates. Inhibitory elbow trajectories were low-pass filtered before plotting and SPM analysis. Significant temporal clusters ($p < 0.05$, two-tailed) are reported with cluster time ranges, durations, and sample sizes. The statistics reflect the results described in the Fig. S3 caption, with normals starting significantly further from the pellet, and excitatories reaching further during the reach phase than inhibitories; and controls being significantly higher than inhibitories briefly at peak reach (t ~ 0.4 sec).

| measure | comparison | n1 | n2 | p | t_start | t_end | duration |
|---|---|---|---|---|---|---|---|
| Elbow X | inhib vs norm | 7 | 4 | 0.004016 | 0.028 | 0.128 | 0.1 |
| Elbow X | excite vs control | 13 | 10 | 0.005017 | 0.252 | 0.38 | 0.128 |
| Elbow X | control vs inhib | 10 | 7 | 0.02056 | 0.38 | 0.444 | 0.064 |
| Elbow X | excite vs inhib | 13 | 7 | 0.03049 | 0.616 | 0.684 | 0.068 |
| Elbow X | excite vs control | 13 | 10 | 0.03922 | 0.512 | 0.552 | 0.04 |
| Elbow X | excite vs norm | 13 | 4 | 1.12e-05 | 0 | 0.272 | 0.272 |
| Elbow X | control vs norm | 10 | 4 | 3.138e-06 | 0 | 0.188 | 0.188 |
| Elbow X | excite vs inhib | 13 | 7 | 8.082e-07 | 0.268 | 0.592 | 0.324 |
| Elbow Y | excite vs norm | 13 | 4 | 0.007382 | 0.364 | 0.548 | 0.184 |

Only significant clusters are shown ($p < 0.05$, two-tailed). n1/n2 denote the number of reaches in each group used for the SPM test; duration = t_end – t_start.

**Table S5.** Significant SPM clusters for shoulder raw coordinate trajectories. Pairwise statistical parametric mapping analyses comparing treatment groups for time-varying shoulder X and Y coordinates. Significant temporal clusters identified by SPM two-sample t-tests ($p < 0.05$, two-tailed) are reported with corresponding time ranges, cluster durations, and sample sizes (number of reaches per group). Only clusters reaching statistical significance are shown.

| measure | comparison | n1 | n2 | p | t_start | t_end | duration |
|---|---|---|---|---|---|---|---|
| Shoulder X | excite vs norm | 13 | 4 | 0 | 0 | 0.712 | 0.712 |
| Shoulder X | inhib vs norm | 7 | 4 | 0.0004476 | 0.44 | 0.58 | 0.14 |
| Shoulder X | excite vs inhib | 13 | 7 | 0.003351 | 0.328 | 0.496 | 0.168 |
| Shoulder X | excite vs control | 13 | 10 | 0.04195 | 0.348 | 0.392 | 0.044 |
| Shoulder X | control vs norm | 10 | 4 | 1.986e-05 | 0.36 | 0.584 | 0.224 |
| Shoulder X | inhib vs norm | 7 | 4 | 4.774e-14 | 0 | 0.336 | 0.336 |
| Shoulder X | control vs norm | 10 | 4 | 5.784e-10 | 0 | 0.344 | 0.344 |
| Shoulder Y | excite vs norm | 13 | 4 | 0.04999 | 0.532 | 0.536 | 0.004 |

Only significant clusters are shown ($p < 0.05$, two-tailed). n1/n2 denote the number of reaches in each group used for the SPM test; duration = t_end – t_start.

**Supplementary Video 1.** Representative rough-and-tumble play bout: a normal cage-mate pinning a lesioned rat. Excerpt from a night-time play recording captured under red-light illumination during the P20–P50 actuation window. The normal animal executes a successful dorsal pin on a lesioned cage-mate, while the lesioned rat attempts to engage with the affected forelimb but is unable to complete a nape grab or maintain a counter-pin, illustrating the forelimb-specific deficit underlying the reduced pin-victory counts in lesioned animals (Fig. 2A).
